# Cell-Surface Proteomic Profiling in the Fly Brain Uncovers New Wiring Regulators

**DOI:** 10.1101/819037

**Authors:** Jiefu Li, Shuo Han, Hongjie Li, Namrata D. Udeshi, Tanya Svinkina, D. R. Mani, Chuanyun Xu, Ricardo Guajardo, Qijing Xie, Tongchao Li, David J. Luginbuhl, Bing Wu, Colleen N. McLaughlin, Anthony Xie, Pornchai Kaewsapsak, Stephen R. Quake, Steven A. Carr, Alice Y. Ting, Liqun Luo

## Abstract

Molecular interactions at the cellular interface mediate organized assembly of single cells into tissues, and thus govern the development and physiology of multicellular organisms. Here, we developed a cell-type-specific, spatiotemporally-resolved approach to profile cell-surface proteomes in intact tissues. Quantitative profiling of cell-surface proteomes of *Drosophila* olfactory projection neurons (PNs) in pupae and adults revealed a global down-regulation of wiring molecules and an up-regulation of synaptic molecules in the transition from developing to mature PNs. A proteome-instructed *in vivo* screen identified 20 new cell-surface molecules regulating neural circuit assembly, many of which belong to evolutionarily conserved protein families not previously linked to neural development. Genetic analysis further revealed that the lipoprotein receptor LRP1 cell-autonomously controls PN dendrite targeting, contributing to the formation of a precise olfactory map. These findings highlight the power of temporally-resolved *in situ* cell-surface proteomic profiling in discovering new regulators of brain wiring.

## INTRODUCTION

In the evolutionary transition from unicellular to multicellular organisms, single cells assemble into highly organized tissues and cooperatively execute physiological functions. To act as an integrated system, individual cells communicate with each other extensively through signaling at the cellular interface. Cell-surface signaling thus controls almost every aspect of the development and physiology of multicellular organisms. Taking the nervous system as an example, cell-surface wiring molecules dictate the precise assembly of the neural network during development (Jan and Jan, 2010; Kolodkin and Tessier-Lavigne, 2011; Sperry, 1963; Zipursky and Sanes, 2010), while neurotransmitter receptors and ion channels mediate synaptic transmission and plasticity in adults (Malenka and Bear, 2004). Delineating the cell-surface signaling is therefore crucial for understanding the organizing principles and operating mechanisms of multicellular systems. Portraying cell-surface proteomes can not only reveal their global landscape and dynamics but also provide a roadmap to investigate the function of individual molecules enriched at specific stages. Cell-surface proteomic profiling has previously been achieved in cultured cells (Loh et al., 2016; Wollscheid et al., 2009). However, cultured cells lose the *in situ* tissue environment that is pivotal for development and physiology *in vivo*.

Here, we present a method for profiling cell-surface proteomes in intact tissues with cell-type and spatiotemporal specificities. Applying this method to *Drosophila* olfactory projection neurons (PNs), we captured cell-surface proteomes of developing and mature PNs and observed globally coordinated dynamics of PN surface proteins corresponding to the wiring and functional stages of the olfactory circuit. A proteomically instructed *in vivo* screen of developmentally enriched PN surface proteins identified 20 new regulators of neural circuit assembly. Further genetic analysis revealed the cell-autonomous function of the evolutionarily conserved lipoprotein receptor LRP1 in dendrite targeting. These data highlight the power of *in situ* cell-surface proteomic profiling in discovering new molecules involved in brain wiring.

## RESULTS

### Biotinylation of PN Surface Proteins in Intact Brains

Despite its functional importance, the cell-surface proteome in intact tissues is difficult to characterize by traditional methods. Biochemical fractionation of membrane proteins not only includes mitochondrial, ER, and Golgi contaminants but also omits secreted and extracellular matrix proteins that form an integral part of the cell-surface proteome (Cordwell and Thingholm, 2010). Moreover, fractionation does not provide cell-type specificity, especially in the nervous system where many cell types intermingle within a compact region. To capture the cell-type-specific, cell-surface proteome in intact tissues, we modified a peroxidase-mediated proximity biotinylation procedure for cultured neurons (Loh et al., 2016) and developed a method for cell-surface biotinylation in intact tissues (Figure 1A; see STAR⋆METHODS for details, including optimized chemical delivery, *in situ* biotinylation reaction, and tissue sample processing). Horseradish peroxidase (HRP) fused to the N-terminus of a generic transmembrane protein, rat CD2, targets HRP to the extracellular side of the plasma membrane (Larsen et al., 2003; Watts et al., 2004). Transgenic expression of HRP-CD2 in genetically defined cell populations thus confers cell-type specificity. Although HRP is cell-surface-targeted, it is also enzymatically active in the secretory pathway intracellularly. To avoid biotinylating proteins there, we used the biotin-xx-phenol (BxxP) substrate, which is impermeable to the plasma membrane (Loh et al., 2016). Combining surface-targeted HRP and membrane-impermeable BxxP, this dual-gate approach should ensure cell-surface specificity. In the presence of hydrogen peroxide (H_2_O_2_), HRP converts BxxP into phenoxyl radicals that promiscuously biotinylate endogenous proteins in proximity. Due to its rapid kinetics, this HRP-mediated reaction requires only a few minutes to complete, providing a cell-surface snapshot with high temporal resolution and minimizing the potential toxicity of H_2_O_2_.

**Figure 1.**
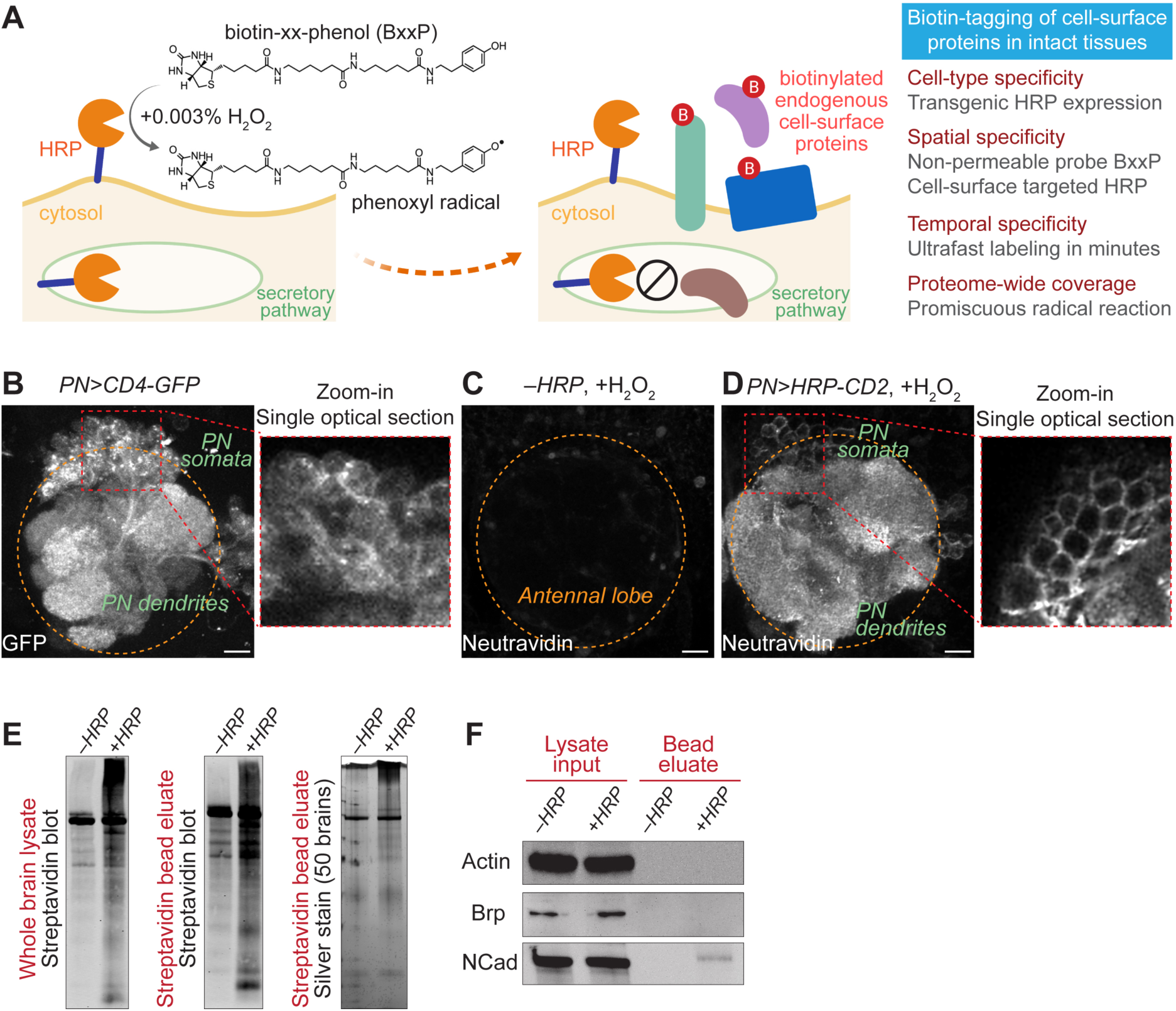
Cell-surface biotinylation of olfactory projection neurons in intact brains. (**A**) Scheme and features of cell-surface biotinylation in intact tissues. (**B**) Olfactory projection neuron (PN) specific *VT033006-GAL4* (*PN-GAL4*, hereafter) drove the expression of membrane-targeted GFP (CD4-GFP). The zoom-in panel shows a single optical section of the PN soma area. Orange circle, antennal lobe. (**C** and **D**) Neutravidin staining of antennal lobes after the cell-surface biotinylation reaction. (C) HRP was not expressed by omitting the *GAL4* driver. (D) *PN-GAL4* drove the expression of cell surface-targeted HRP (HRP-CD2). The zoom-in panel shows a single optical section of the PN soma area. (**E**) Left and middle, streptavidin blots of the whole-brain lysate (left) and the post-enrichment bead eluate (middle). Right, silver stain of the post-enrichment bead eluate (right). *–HRP*, *PN-GAL4* omitted; *+HRP*, *PN>HRP-CD2*. (**F**) Immunoblots of intracellular proteins, actin and bruchpilot (Brp), and neuronal surface protein N-cadherin (NCad) in the whole-brain lysate and the post-enrichment bead eluate. *–HRP*, *PN-GAL4* omitted; *+HRP*, *PN>HRP-CD2*. Scale bar, 10 μm. See also Figure S1.

We applied this cell-surface biotinylation strategy to *Drosophila* olfactory projection neurons (PNs). In the antennal lobe, the first-order olfactory processing center of *Drosophila*, PNs of the same type target their dendrites to a specific glomerulus to form synaptic connections with their partner olfactory receptor neurons (ORNs) (Figures 1B and S1A). Precise one-to-one pairings of 50 types of PNs and ORNs at 50 discrete glomeruli build an anatomically stereotyped olfactory map in the antennal lobe (Benton et al., 2009; Couto et al., 2005; Fishilevich and Vosshall, 2005; Gao et al., 2000; Jefferis et al., 2001; Silbering et al., 2011; Vosshall and Stocker, 2007; Vosshall et al., 2000), providing an excellent system for studying neural development and computation (Hong and Luo, 2014; Wilson, 2013). To examine HRP-mediated cell-surface biotinylation in intact brains, we first stained adult brains with fluorophore-conjugated neutravidin, which specifically recognizes biotin. We observed extensive, HRP-dependent biotinylation of PN dendrites and axons (Figures 1C, 1D, S1B, and S1C). Confocal optical sectioning revealed that only the surface of PN cell bodies was biotinylated, in contrast to the broad distribution of membrane-targeted GFP (zoom-in panels of Figures 1B and 1D).

Biochemical characterization by streptavidin blot and silver stain of the brain lysate or its post-enrichment eluate showed that the PN surface reaction biotinylated a wide range of proteins compared with the control lacking HRP-CD2 expression, although fly brains also expressed endogenously biotinylated proteins that require filtering in proteomic analysis (Figure 1E). Immunoblotting revealed that the neuronal surface marker N-cadherin, but not the cytosolic proteins actin or bruchpilot (a pan-neural synaptic marker), was detected in streptavidin bead eluates in an HRP-dependent manner (Figure 1F). Thus, our surface biotinylation procedure provides a way to label and enrich cell-surface proteins of a chosen cell type in intact tissues, enabling mass spectrometry-based proteomic profiling.

### Cell-Surface Proteomes of Developing and Mature PNs

It is generally assumed that cell-surface proteins would change as neurons mature, but this has not been systematically examined due to technical limitations. Here, we profiled the PN surface proteome at two time points – 36 hours after puparium formation (36hAPF) when developing PNs elaborate their dendrites and axons to build synaptic connections (Jefferis et al., 2004), and 5 days (5d) after eclosion into adults when mature PNs actively process olfactory information (Wilson, 2013) (Figure 2A). To better quantify protein dynamics and filter out contaminants captured in negative controls, we used an 8-plex tandem mass tag (TMT)-based quantitative strategy (Thompson et al., 2003) and profiled each time point with two biological replicates (Figure 2B). Each time point also contained two negative controls (Figure 2B; without HRP-CD2 expression or omitting H_2_O_2_ in the reaction) that capture endogenously biotinylated proteins and non-specific binders to streptavidin beads. Freshly dissected brains (∼1000 per TMT plex) were incubated with the BxxP substrate for one hour before a 5-minute H_2_O_2_ reaction. Biotinylated proteins were then enriched from brain lysates using streptavidin beads, followed by on-bead trypsin digestion, TMT labeling, and liquid chromatography-tandem mass spectrometry analysis (LC-MS/MS) (Figure 2A).

**Figure 2.**
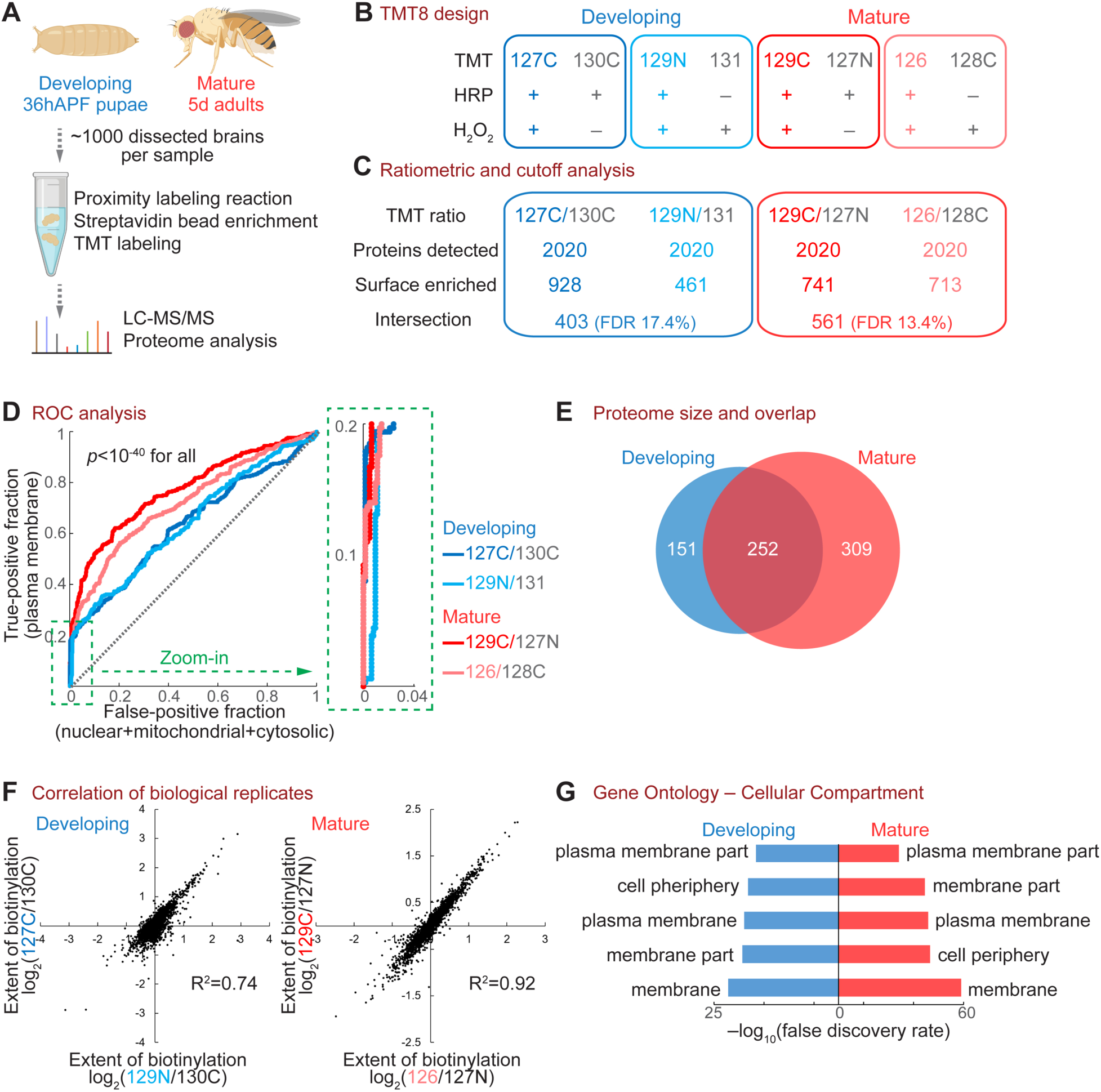
Cell-surface proteomic profiling of developing and mature PNs. (**A**) Workflow of the PN surface proteomic profiling. (**B**) Design of the 8-plex tandem mass tag (TMT8)-based quantitative proteomic experiment. Each time point comprised two biological replicates (blue or red) and two negative controls (grey). (**C**) Numbers of proteins after each step of the ratiometric and cutoff analysis. (**D**) Receiver operating characteristic (ROC) curve of each biological replicate. Proteins were ranked in a descending order based on the TMT ratio. True-positive denotes plasma membrane proteins curated by the UniProt database. False-positive includes nuclear, mitochondrial, and cytosolic proteins without membrane annotation by UniProt. The top 20% region (dotted green box) is enlarged on the right. A two-sample Wilcoxon rank-sum test was used to determine the statistical significance. (**E**) Venn diagram of developing and mature PN surface proteomes, after ratiometric and cutoff analyses. (**F**) Correlation of biological replicates. (**G**) Top 5 Cellular Compartment terms of the developing and mature PN surface proteomes in gene ontology analysis. See also Figure S2.

From this experiment, 2020 proteins were detected with 2 or more unique peptides (Figure 2C). To remove potential contaminants, we adopted a ratiometric strategy (Hung et al., 2014) in which the TMT ratio of each protein reflects its differential enrichment in the reaction group and the negative control group (Figure 2C). A *bona fide* PN surface protein should exhibit a high TMT ratio, as it should be extensively biotinylated in the reaction group but not in the control group. A false positive, such as an endogenously biotinylated protein or a non-specific bead binder, would have a low ratio, as it should be captured similarly in both groups. We ranked each biological replicate in a descending order by the TMT ratio and plotted its receiver operating characteristic (ROC) curve (Figure 2D), which depict the true-positive rate against the false-positive rate of detected proteins. Notably, the top 20% of proteins exhibited almost vertical ROC curves (zoom-in in Figure 2D), demonstrating high specificity. To maximize the signal-to-noise ratio of the proteome, we cut off each biological replicate at the position where the value of *true-positive rate – false-positive rate* maximized (Figure S2A) and collected only the overlapping proteins of the two replicates at each time point to further minimize potential contaminants (Figure 2C).

Through ratiometric and cutoff analyses, we obtained developing and mature PN surface proteomes containing 403 and 561 proteins, respectively (Figures 2C and 2E). Both proteomes exhibited high correlations between biological replicates (Figures 2F and S2B) and high spatial specificity as seen from the cellular compartment classification in gene ontology (Figure 2G), in line with our histological and biochemical characterizations (Figures 1D and 1F).

### Temporal Evolution of the PN Surface Proteome

Of the 712 proteins in the PN surface proteomes, only 252 proteins were shared by the developing and mature proteomes, whereas 460 were specific to either developing or mature PNs (Figure 2E). These data suggest a profound difference between the PN surface proteomes at these time points. We hence systematically examined and compared the developing and mature PN surface.

In contrast to their nearly identical cell-surface annotations (Figure 2G), the developing and mature proteomes exhibited distinct and non-overlapping signatures regarding their biological functions (Figure 3A). In accord with the primary task of PNs at each time point, the developing surface highlighted the processes for neural development while the mature surface featured categories covering ion channels, receptors, and transporters (Figure 3A). The most enriched proteins from developing PN surface with known functions were predominantly neural development molecules, in particular those involved in axon guidance and neural circuit wiring (Figure 3B and Table 1). This is surprising, as guidance molecules are generally thought to act at growth cones – only the tip of arborizing dendrites and axons. Strikingly, the top 100 cell-surface enriched proteins of developing PNs contained almost all cell-surface regulators of olfactory circuit wiring identified in the past two decades, as well as many wiring molecules discovered in other systems (Table 1). By contrast, the most enriched proteins of mature PN surface with known functions were not synaptic molecules (Figure 3B), consistent with the notion that synapses are highly restricted at select loci of a mature neuron. The enrichment of many previously uncharacterized proteins on the mature PN surface (Figure 3B) suggests that future investigations of these proteins may reveal new facets of PN physiology.

**Figure 3.**
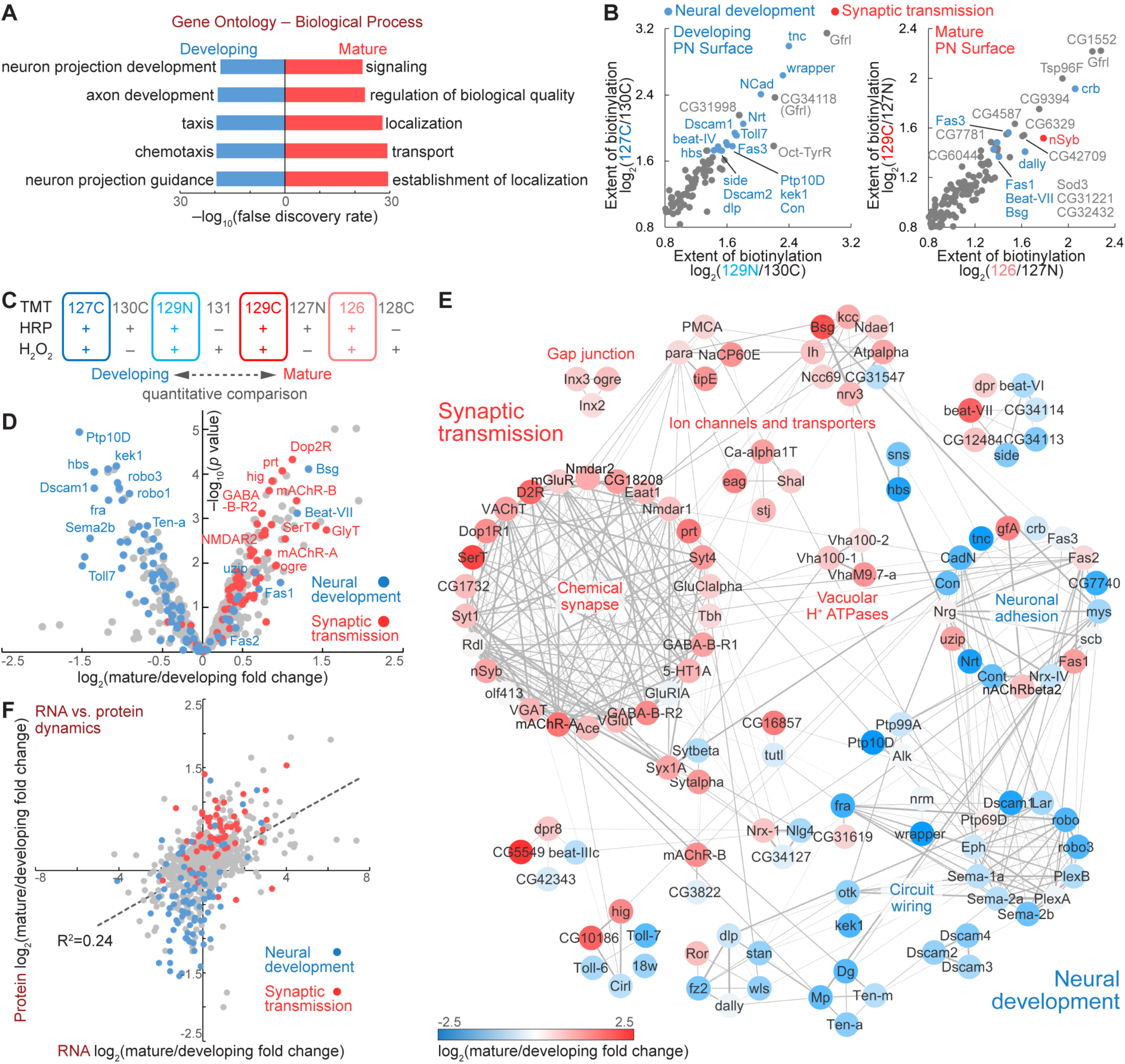
Temporal evolution of the cell-surface proteome in accord with PN development and function. (**A**) Top 5 Biological Process terms of the developing and mature PN surface proteomes in gene ontology analysis. (**B**) Most enriched proteins on the developing and mature PN surface, respectively. Blue, known neural development molecules; red, known synaptic transmission molecules. (**C** and **D**) TMT-based quantification revealed the expression dynamics of cell-surface proteins in PN development and maturation. (**E**) Coordinated dynamics of functionally associated molecular complexes in developing and mature PNs. Markov clustering was performed with protein-protein interaction scores from the STRING database. Grey lines denote protein-protein interactions. Nodes are color-coded based on the expression dynamics as in Figure 3D. (**F**) RNA vs. protein dynamics of PN surface molecules. Dashed line depicts the linear regression. See also Figure S3.

**Table 1.**
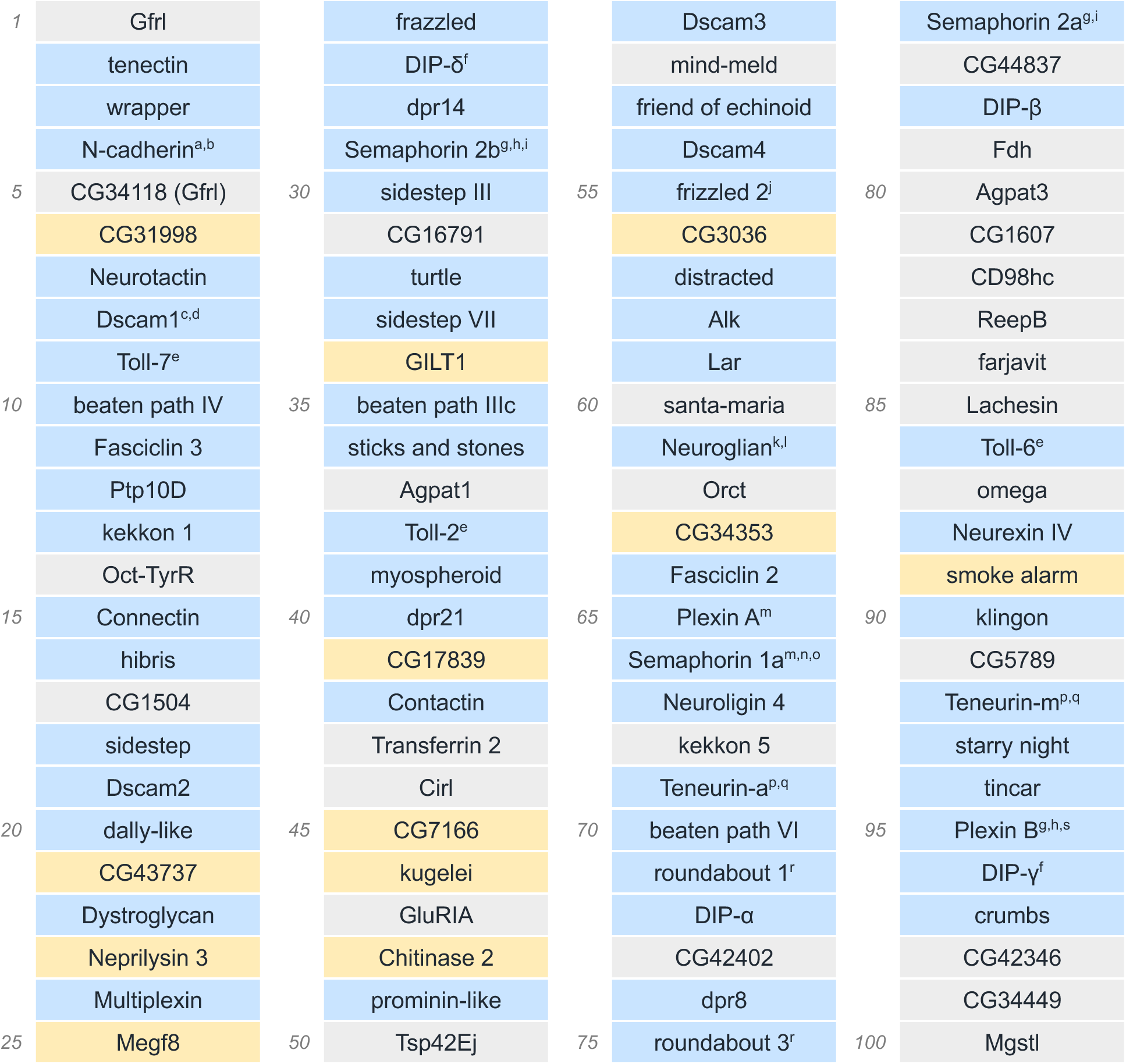
Top 100 cell-surface enriched proteins in developing PNs. Proteins are ranked by the 127C:130C TMT ratio (see Figure 2B). Blue background denotes proteins with known function in neural development, in particular circuit wiring. Yellow denotes newly identified wiring molecules in this study (Figure 4 and Table 2). References for previously identified wiring molecules in the fly olfactory circuit are as follows: ^a^Hummel and Zipursky, 2004; ^b^Zhu and Luo, 2004; ^c^Hummel et al., 2003; ^d^Zhu et al., 2006; ^e^Ward et al., 2015; ^f^Barish et al., 2018; ^g^Joo et al., 2013; ^h^Li et al., 2018; ^i^Sweeney et al., 2011; ^j^Singh et al., 2010; ^k^Chen and Hing, 2008; ^l^Kaur et al., 2019; ^m^Sweeney et al., 2007; ^n^Komiyama et al., 2007; °Shen et al., 2017; ^p^Hong et al., 2012; ^q^Mosca and Luo, 2014; ^r^Jhaveri et al., 2004; and ^s^Guajardo et al., 2019.

Besides examining the developing and mature proteomes separately (Figures 3A and 3B), the TMT-based profiling allowed us to examine the quantitative dynamics of each individual protein (Figure 3C). We found that most known neural development and synaptic transmission molecules segregated from each other, falling into the left and right quadrants, respectively (Figure 3D), revealing a global down-regulation of wiring molecules and an up-regulation of synaptic molecules in the transition from developing to mature PNs. Next, we clustered the PN surface proteins with known neural development or synaptic functions by their protein-protein interactions (Figure 3E) and observed core molecular complexes for neural development (e.g., ‘circuit wiring’) and physiology (e.g., ‘chemical synapse’ and ‘gap junction’). Most clusters comprised proteins with unidirectional dynamics, as seen by separately clustered blue or red nodes (Figure 3E), revealing the concerted regulation of functionally associated proteins in developing and mature neurons. Interestingly, the cluster of neuronal adhesion molecules showed mixed temporal changes (‘neuronal adhesion’ in Figure 3E). This is consistent with their biological functions: cell adhesion proteins play central roles in circuit assembly at the developmental stage and continue to maintain inter-cellular adhesion throughout the lifetime. Indeed, some adhesion proteins were among the most enriched proteins of the mature PN surface (Figure 3B), and were even up-regulated in mature PNs (blue dots in the right quadrant of Figure 3D).

To compare the RNA and protein dynamics of PN surface molecules in the developing-to-mature transition, we also profiled the PN transcriptomes from 36hAPF pupae and 5d adults (Figure S3A–S3D). While the transcriptomic dynamics predicted the global trend of proteomic dynamics, it failed to forecast the temporal change of many individual molecules (Figure 3F), similar to the previous observation by comparing whole-cell proteomes and transcriptomes of brains (Carlyle et al., 2017). For example, the static mRNA level did not predict the differential expression of many proteins involved in synaptic transmission and neural development (red and blue dots near the y-axis of Figure 3F). Moreover, some functionally important molecules exhibited inverse dynamics by RNA and protein. In most cases, the protein dynamics better agreed with the function of that molecule (Figures S3E and S3F). Many factors could contribute to the discrepancy between RNA and protein dynamics, including post-transcriptional regulation, temporal lag of translation, and cell-surface delivery of proteins.

The discrepancy between RNA and protein dynamics could also be contributed by the fact that PN surface environment is not solely determined by PN themselves but also contributed by nearby cells. The radical-mediated proximity biotinylation (Figure 1A) should survey the entire PN surface environment including secreted and transmembrane proteins produced by adjacent cells. Indeed, by comparing the cell-surface proteomes with the transcriptomes of PNs, we observed a small fraction of molecules captured on the PN surface with very low read counts in PN RNA-sequencing (Figure S3G). These molecules are likely expressed by nearby non-PN cells, although we cannot exclude the possibility that a low amount of stable RNA in PNs produces plentiful proteins.

### Unconventional Wiring Molecules Identified by Proteome-Instructed RNAi Screen

Many proteins previously identified for their function in regulating wiring specificity of PNs and other neurons exhibited developmentally enriched expression (blue dots in the left quadrant of Figure 3D). This raised the question of whether the previously uncharacterized proteins enriched in developing PN surface (grey dots in the same quadrant) could play similar functions. We thus carried out a genetic screen of these novel, developmentally enriched PN surface proteins based purely on their proteomic profile (Figure 4A) to test whether they participate in neural circuit assembly. We previously developed an *in vivo* RNAi-based genetic screen (Xie et al., 2019) featuring pan-neuronal knockdown of candidate molecules and simultaneous monitoring of the dendrite and axon targeting specificity of two types of PNs and two types of ORNs, respectively, in the ventromedial (VM) antennal lobe (Figure 4B). In wild-type flies, these dendrites and axons target to stereotyped locations in the antennal lobe with high precision (Figure 4C), enabling a confocal-based high-resolution screen for PN dendrite targeting, ORN axon targeting, and PN-ORN synaptic partner matching.

**Figure 4.**
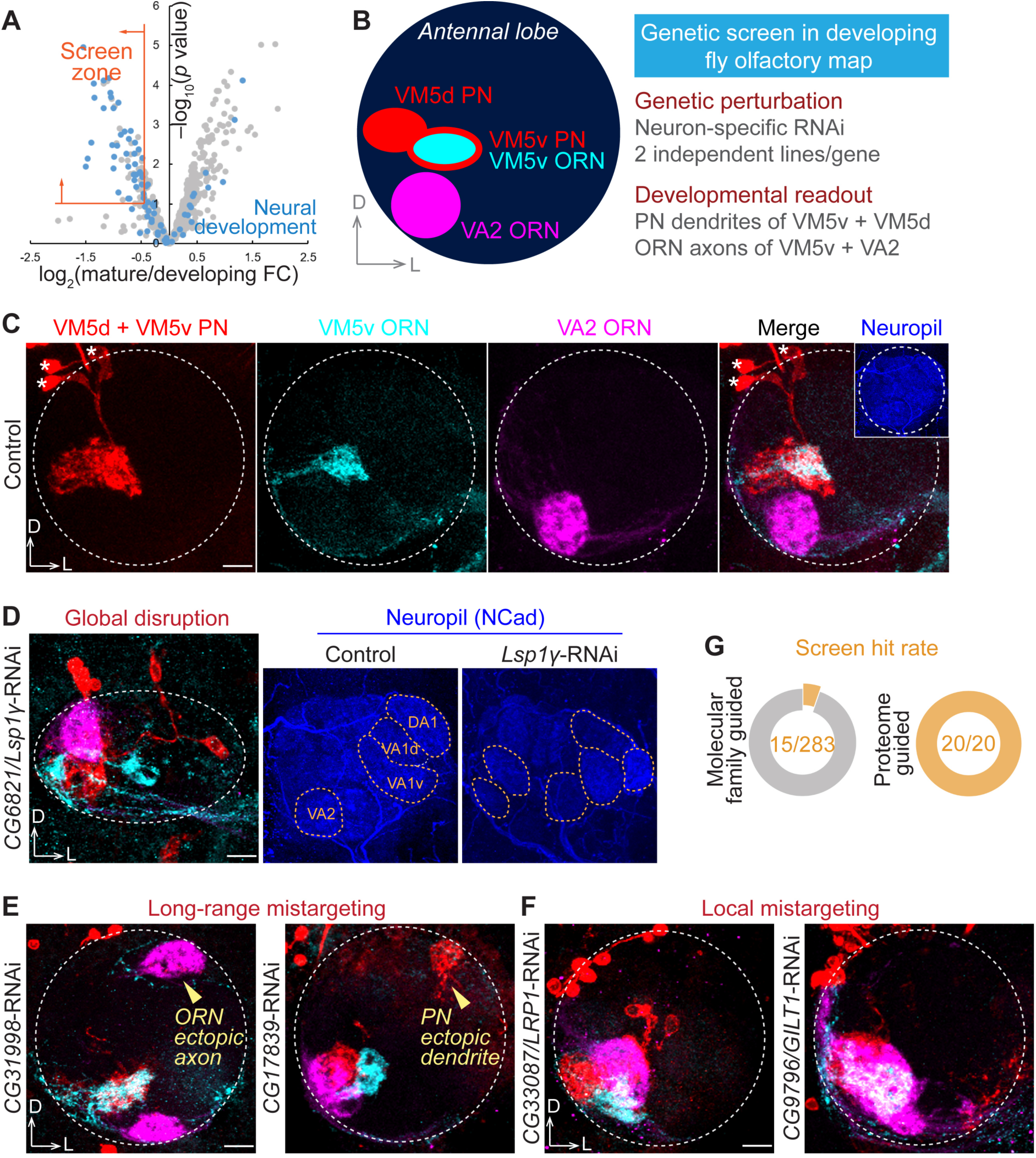
Genetic screen identified new regulators of neural circuit assembly. (**A**) Selection criteria for the genetic screen. Screen zone cutoffs: log_2_(mature/developing FC) < – 0.4 and –log_10_(*p* value) > 1. FC, fold change. (**B**) Scheme and features of the genetic screen in ventromedial (VM) PNs and ORNs. VM5d and VM5v PNs were labelled by *GMR86C10-LexA*-driven membrane-targeted tdTomato (*LexAop-mtdTomato*; red). VM5v and VA2 ORNs were labelled by *Or98a* promoter-driven membrane-targeted GFP (*Or98a-mCD8-GFP*; cyan) and *Or92a* promoter-driven rat CD2 transmembrane motif (*Or92a-rCD2*; magenta), respectively. The pan-neuronal *C155-GAL4* drove the expression of gene-specific RNAi. (**C**) Targeting of VM5d/v PN dendrites and VM5v/VA2 ORN axons in a control antennal lobe. None of the 62 examined antennal lobes in controls exhibited any targeting defects. Dashed circle, antennal lobe; asterisk, PN soma. (**D**) *CG6821/Lsp1γ* knockdown caused global disruption of the antennal lobe structure. Yellow dashed circles in neuropil staining (blue) represent stereotyped glomeruli that are easily identified in control but misshapen and unrecognizable in *Lsp1γ* knockdown. (**E**) *CG31998* and *CG17839* knockdown caused long-range mistargeting of ORN axons and PN dendrites, respectively, to the dorsolateral (DL) antennal lobe. (**F**) *CG33087/LRP1* and *CG9796/GILT1* knockdown caused VM local mistargeting of ORN axons and PN dendrites. (**G**) Hit rates of our previous molecular family-based screen (Xie et al., 2019) and the current cell-surface proteome guided screen, with the identical assay (scheme in B). Scale bar, 10 μm. D, dorsal; L, lateral. RNAi phenotypic penetrances are listed in Table 2. See also Figure S4.

**Table 2.**
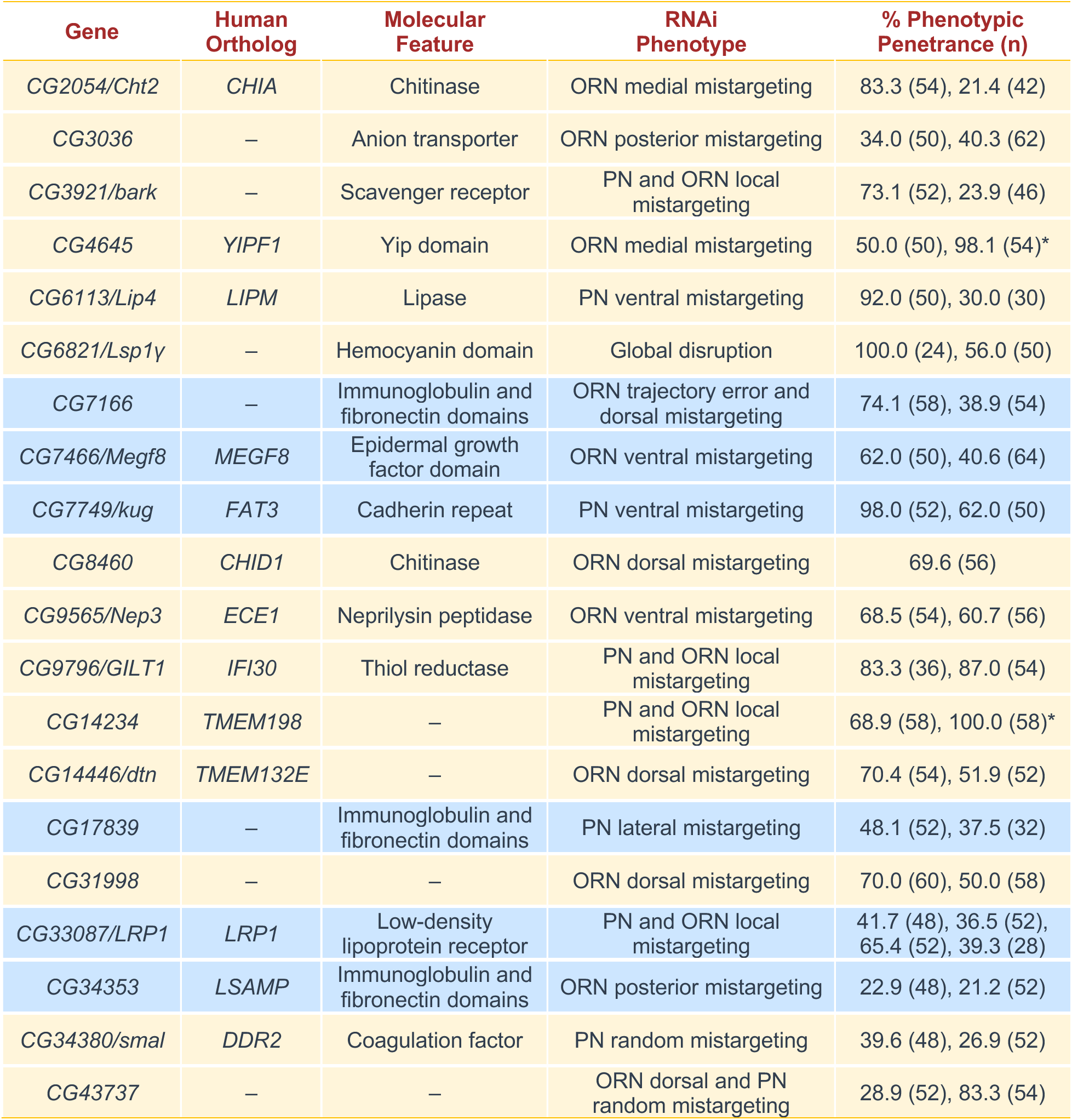
RNAi screen in ventromedial (VM) PNs and ORNs: genes, molecular features, phenotypes, and penetrances. Human orthologs were searched by the FlyBase Homologs Search tool. Only the orthologs consistently identified by four or more databases are listed. Molecular features were searched through FlyBase and UniProt. Blue background denotes proteins belonging to families of classic wiring molecules based on their structural domains; yellow denotes proteins from molecular families not previously linked to neural development. Phenotypic penetrance of each RNAi is listed with the number of antennal lobes examined in parentheses. Antennal lobe image of each RNAi is included in Figure S4. Asterisks represent two cases where pan-neuronal RNAi was lethal and *PN-GAL4* was used instead to drive RNAi expression.

We observed diverse and distinct wiring defects in all 20 genes we screened (Figures 4D–4F and Table 2). In 19 of the 20 genes, consistent phenotypes were observed using at least two independent RNAi lines targeting non-overlapping regions (Figure S4). For example, knocking down the secreted protein Lsp1γ (larval serum protein 1 γ) caused global disruption of the antennal lobe, in which the glomerular identities were not recognizable in neuropil staining (Figure 4D). Consequently, the targeting of VM PNs and ORNs was randomized (Figure 4D). Unlike Lsp1γ, disruption of other molecules led to more specific targeting defects. RNAi against two previously uncharacterized molecules, CG31998 and CG17839, caused long-range mistargeting of ORN axons and PN dendrites, respectively, to the dorsolateral (DL) antennal lobe (Figure 4E), suggesting that they may participate in coarse organization of the olfactory map, like Semaphorins and their receptors (Joo et al., 2013; Komiyama et al., 2007; Li et al., 2018; Sweeney et al., 2011). In contrast, knocking down LRP1 (low density lipoprotein receptor protein 1) and GILT1 (gamma-interferon-inducible lysosomal thiol reductase 1, a predicted cell-surface protein (Almagro Armenteros et al., 2017; Tan et al., 2009) with ‘lysosomal’ in its name due to phylogenetic homology) led to local mistargeting defects confined within the VM antennal lobe (Figure 4F), indicating that these molecules likely regulate local refinement in target selection, resembling Teneurins and two leucine-rich repeat proteins – Capricious and Tartan (Hong et al., 2009, 2012).

Many known wiring molecules carry common structural domains, such as cadherin repeat, immunoglobulin domain, and leucine-rich repeat, which often mediate the molecular interactions required for neuronal recognition (de Wit et al., 2011; Kolodkin and Tessier-Lavigne, 2011). In our previous RNAi screen using the same strategy (Xie et al., 2019) (Figure 4B), we selected candidates containing those structural signatures and identified 15 putative regulators of olfactory wiring out of 283 screened candidates, with a hit rate of ∼5% (Figure 4G). Here, we screened 20 developmentally enriched PN surface proteins without considering their molecular families (Figure 4A) and achieved a striking hit rate of 100% (Figure 4G). Notably, only a small fraction of them shared the structural signatures of classic wiring molecules described above. Most came from molecular families that had not previously been associated with neural development (Table 2). 13 of these 20 proteins have clear human orthologs, suggesting that our knowledge of molecules involved in assembling neural circuits, whether in flies or in mammals, is far from complete.

### LRP1 Cell-Autonomously Regulates PN Dendrite Targeting

From the screen hits, we carried out an in-depth analysis of LRP1, an evolutionarily conserved protein (Figure 5A) involved in the pathogenesis of atherosclerosis and Alzheimer’s disease (Boucher et al., 2003; Kanekiyo et al., 2013; Kang et al., 2000; Liu et al., 2017; Tachibana et al., 2019). Although loss of LRP1 has been shown to impair motor behavior (May et al., 2004), brain insulin signaling (Brankatschk et al., 2014), and growth cone dynamics *in vitro* (Landowski et al., 2016), whether it regulates neural development *in vivo* was unknown.

**Figure 5.**
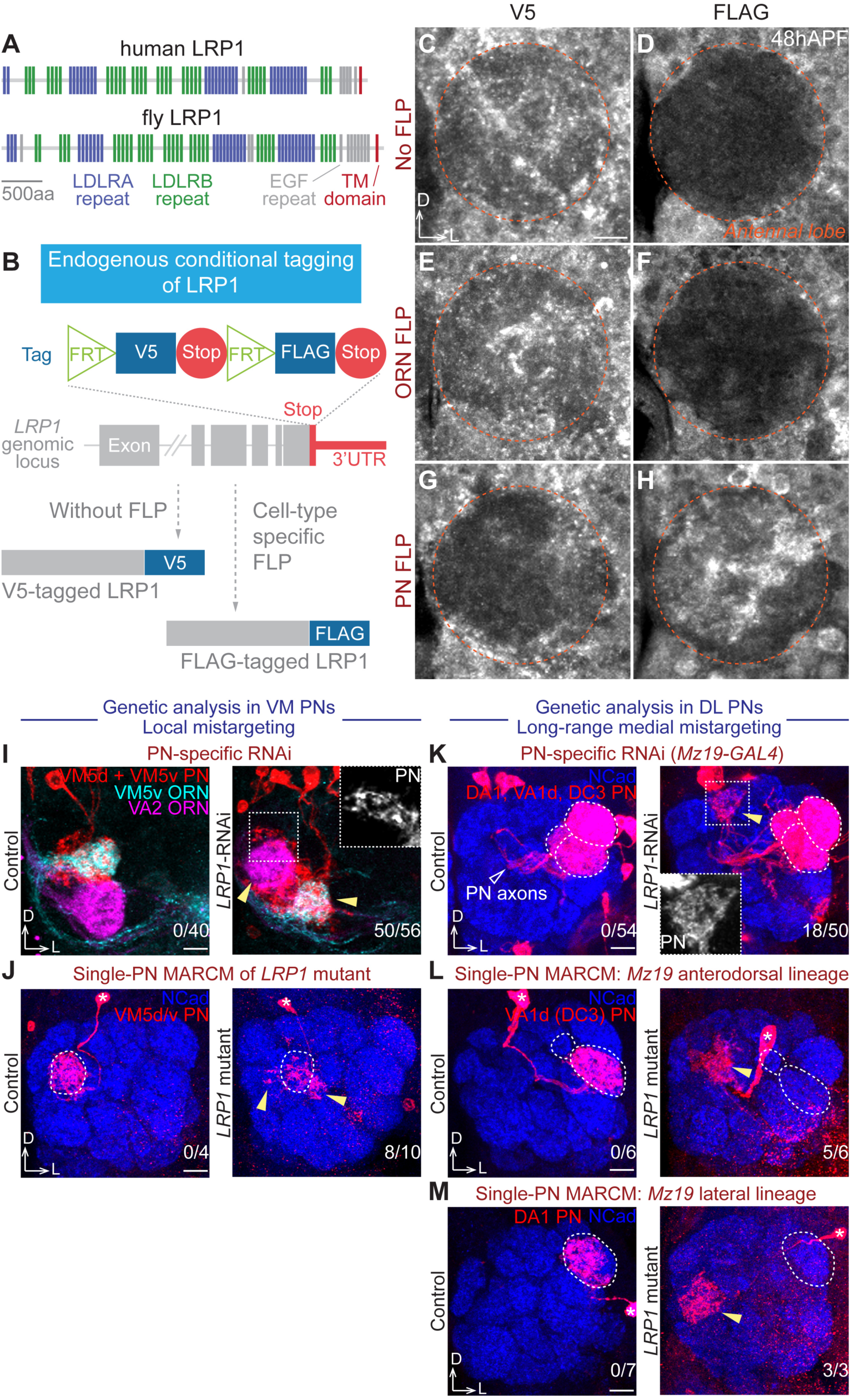
LRP1 cell-autonomously controls PN dendrite targeting. (**A**) Domain structures of human and fruit fly LRP1 proteins. TM, transmembrane. (**B**) Schematic of conditional tagging of LRP1 to reveal its cell-type-specific endogenous protein expression pattern. (**C**–**H**) V5 (C, E, G) or FLAG (D, F, H) staining of tagged LRP1 under different conditions: without FLP (C and D), ORN-specific *eyFLP* (E and F), and PN-specific *VT033006>FLP* (G and H). 48hAPF, 48 hours after puparium formation. Orange circle, antennal lobe. Cortex glia outside of the antennal lobe have high background signal in FLAG staining. (**I**) PN-specific *LRP1*-RNAi knockdown caused local mistargeting of VM5d/v PN dendrites and VM5v/VA2 ORN axons (zoom-in square and arrowheads). (**J**) Mosaic analysis of *LRP1* mutant in single VM5d/v PNs showed dendrite local mistargeting (arrowheads). Dotted circle, normal targeting area. (**K**) PN-specific *LRP1*-RNAi knockdown caused long-range medial mistargeting (zoom-in square and yellow arrowhead) of *Mz19*+ PN dendrites, which normally target the dorsolateral (DL) DA1, VA1d, and DC3 glomeruli (dotted circles). (**L** and **M**) Mosaic analysis of *LRP1* mutant in single *Mz19+* PNs showed long-range medial dendrite mistargeting (arrowheads). Dotted circle, normal targeting area. (L) *Mz19+* anterodorsal lineage PNs, which normally target dendrites to VA1d or DC3. (M) *Mz19+* lateral lineage PNs, which normally target dendrites to DA1. Scale bar, 10 μm. D, dorsal; L, lateral. Asterisk, PN soma. The number of antennal lobes with mistargeting over the total number of antennal lobes examined is noted at the bottom-right corner of each panel. See also Figure S5 and Figure S6.

Pan-neuronal RNAi knockdown of LRP1 caused local mistargeting of both PN dendrites and ORN axons in the VM antennal lobe (Figures 4F and S4Q), raising the question of whether LRP1 functions in PNs, ORNs, olfactory local interneurons, or any of these combinations. Antibody staining showed that LRP1 proteins were expressed in the antennal lobe at 48hAPF, when PN dendrites and ORN axons interact to establish targeting specificity (Figure S5A), but could not provide information about its cellular origin. To address the question of cellular origin, we devised a CRISPR-based conditional tagging strategy, in which a FLP-gated protein tag cassette replaced the endogenous stop codon of *LRP1* without removing any coding or regulatory sequences (Figure 5B). Indeed, tagged *LRP1* completely rescued the lethality of *LRP1* mutant, indicating no functional disruption caused by tagging. Consistent with the tag design (Figure 5B), no FLAG signal was detected in the antennal lobe in the absence of FLP expression (Figure 5D), while the V5 staining resembled LRP1 antibody staining (Figures 5C and S5A). With ORN-specific FLP, there was still no FLAG signal (Figure 5F), indicating that LRP1 is not expressed in ORN axons. By contrast, PN-specific FLP resulted in strong FLAG signal in the antennal lobe (Figure 5H), along with an almost complete loss of V5 staining (Figure 5G), revealing that LRP1 is predominately expressed in PN dendrites. Consistently, our single-cell RNA sequencing (Li et al., 2017, 2019) also found that *LRP1* is expressed in PNs but not ORNs (Figure S5C).

To test the functional requirement of LRP1 in specific olfactory neuron types, we performed cell-type-specific RNAi knockdown and MARCM-based mosaic analysis (Lee and Luo, 1999) (Figure S6C) using a null mutant (Figures S6A and S6B). Consistent with the expression pattern, ORN-specific RNAi knockdown did not cause any wiring defects (Figure S6D). In *eyFLP*-based MARCM, *eyFLP* expression restricted *LRP1* loss to ORNs, and only homozygous mutant ORNs expressed the fluorescent marker GFP while the other wild-type or heterozygous cells were not labelled. In line with the expression and ORN-specific knockdown, loss of *LRP1* in ORNs did not cause any targeting defects (Figures S6E and S6F). Thus, the ORN axon mistargeting phenotype was likely caused by loss of *LRP1* in PNs.

Indeed, PN-specific knockdown (Figure 5I) phenocopied pan-neuronal knockdown (Figure 4F), resulting in both PN dendrite and ORN axon mistargeting in the VM antennal lobe. Thus, the ORN phenotype was a non-autonomous effect caused by loss of LRP1 in PN dendrites. In *hsFLP*- and *GMR86C10-GAL4*-based MARCM analysis, single-cell clones of *LRP1^™/™^* VM5d or VM5v PNs extended dendrites to nearby glomeruli outside of the VM5d/v glomeruli (Figure 5J), recapitulating the RNAi knockdown phenotype and demonstrating that LRP1 acts cell-autonomously in VM5d/v PN dendrite targeting.

To test whether LRP1 is also required in dendrite targeting of other PN types, we used *Mz19-GAL4*, which labels DA1 PNs of the lateral neuroblast lineage and VA1d/DC3 PNs of the anterodorsal neuroblast lineage, all of which target dendrites to the dorsolateral (DL) antennal lobe (Jefferis et al., 2001). In contrast to the local defects in VM PNs, we observed long-range medial mistargeting when knocking down LRP1 in *Mz19*+ PNs (Figure 5K). In *Mz19-GAL4*-based MARCM, single-cell clones of *LRP1^™/™^* PNs from both lineages showed mistargeting to the medial antennal lobe (Figures 5L and 5M). Taken together, our genetic analyses revealed that LRP1 cell-autonomously regulates different aspects of dendrite targeting in distinct PN populations – local refinement of VM PNs and long-range targeting of DL PNs.

## DISCUSSION

Systematic characterizations of cell-surface proteomes should facilitate the understanding of cell-cell communication in development and physiology of multicellular organisms. We describe here, for the first time to our knowledge, a method for profiling cell-surface proteomes in intact tissues with cell-type and spatiotemporal specificities. Applying it to *Drosophila* olfactory projection neurons enabled us to reveal systematic changes of cell-surface proteomes in developing and mature neurons, and identify new classes of evolutionarily conserved wiring molecules. We discuss below these technological and biological advances.

### Profiling Cell-Surface Proteomes and Their Dynamics in Intact Tissues

Mass spectrometry-based proteomics provides a systemic way for understanding proteome and its dynamics in biological systems, including the nervous system (Aebersold and Mann, 2016; Han et al., 2018; Hosp and Mann, 2017; Natividad et al., 2018). Despite its central roles in neural development and function, the cell-surface proteome of a specific neuronal type in intact brains was difficult to characterize due to the lack of an appropriate method. Chemical labeling enables the enrichment of cell-surface proteins (Nunomura et al., 2005; Wollscheid et al., 2009; Zhang et al., 2003) but does not provide cell-type specificity. Alternative technologies for cell-type-specific proteomics, such as bio-orthogonal metabolic labeling (Alvarez-Castelao et al., 2017; Liu et al., 2018) or organelle tagging (Fecher et al., 2019), are not amenable to the analysis of cell-surface proteomes.

Here, we report a spatiotemporally-resolved approach to profile the cell-surface proteome of a genetically defined cell population in intact tissues. Compared with the previous methods for profiling the cell-surface molecular composition, our approach simultaneously enables cell-type, subcellular, and temporal specificities in an *in situ* tissue context. Our quantitative profiling of PNs not only provides a detailed view of neuronal cell-surface proteomes at the wiring and functional stages (Figure 3B), but also systematically uncovers the dynamics of individual proteins across these two stages (Figure 3D). We found that the cell surface of developing PNs is highly enriched for wiring molecules (Table 1), and there is a proteome-wide coordinated dynamics of PN surface molecules during the transition from developing to mature stages (Figure 3E).

Our approach should be readily applicable for profiling proteins on the surface of other cell types, tissues, and organisms by expression of cell-surface delivered HRP from a transgene in desired cell type(s). In addition to studying cell-cell communications under physiological conditions, *in situ* quantitative cell-surface proteomic profiling can be used to decipher proteomic changes in mutants or under pathological conditions. Cell-surface proteins are also the main targets for drug development (Christopoulos, 2002). Probing cell-surface proteome changes under pathogenic conditions and in response to drug application can help identify therapeutic targets and monitor treatment efficacy.

### Identification of Unconventional New Wiring Molecules

In the past decades, identification and mechanistic studies of classic wiring molecules have revealed many fundamental principles governing neural circuit assembly (Jan and Jan, 2010; Kolodkin and Tessier-Lavigne, 2011; Zipursky and Sanes, 2010). Despite these advances, our current knowledge is far from explaining the striking precision of connectivity observed in the nervous system. Remarkably, our cell-surface proteomic profiling enriched almost all olfactory wiring regulators identified in the past two decades (Table 1). This inspired us to perform a proteome-instructed, unbiased screen for previously uncharacterized molecules. Indeed, this screen was exceptionally efficient (Figure 4G) in uncovering novel cell-surface molecules controlling circuit assembly (Table 2). The discovery of LRP1 as a cell-autonomous regulator of dendrite targeting was surprising, as it has been extensively studied in the nervous system for its involvement in Alzheimer’s disease (Kanekiyo et al., 2013; Kang et al., 2000; Liu et al., 2017; Tachibana et al., 2019) but no *in vivo* neurodevelopmental function was previously reported. Given its widely observed role as an endocytic receptor (Herz and Strickland, 2001; Rohlmann et al., 1998), it is likely that LRP1 controls via endocytosis the dynamics of specific ligands or receptors in distinct PN types, thereby accounting for cell-type-specific and stereotyped loss-of-function wiring defects (Figures 5I–5M).

As summarized in Table 2, the precision of neural circuit assembly requires multiple previously unexpected classes of proteins that are conserved from flies to humans. Since many of these proteins would not be identified by a molecular family-based screen while a genome-wide unbiased screen *in vivo* is technically challenging, our study highlights the power of temporally-resolved cell-surface proteomic profiling in discovering novel regulators of brain wiring. Future investigations of these proteins should expand our understanding of the mechanisms by which neural circuits are assembled with exquisite specificity.

## ACKNOWLEDGEMENTS

We thank Suzanne Eaton for her kind gifts of LRP1 reagents and mourn her tragic death in July 2019. We thank Y. Aso, the Bloomington *Drosophila* Stock Center, the Vienna *Drosophila* Resource Center, the Kyoto *Drosophila* Genomics and Genetic Resources Center, and the Japan National Institute of Genetics for fly lines; T. Clandinin, X. Gao, K. C. Garcia, Y. Ge, J. Kebschull, T. Mosca, D. Pederick, J. Ren, K. Shen, A. Shuster, M. Simon, Y. Takeo, D. Wang, B. Zhao, and members of the Luo and Ting laboratories for technical support, advice, and/or comments on this study; M. Molacavage and S. Wheaton for administrative assistance. J.L. and L.L. thank The Yingwei Cui and Hui Zhao Neuroscience Fund for support. J.L. was supported by a Genentech Foundation predoctoral fellowship and a Vanessa Kong-Kerzner graduate fellowship. S.H. is supported by a Stanford Bio-X Bowes graduate fellowship. H.L. was a Wu Tsai Neurosciences Institute interdisciplinary scholar. A.Y.T. is an investigator of the Chan Zuckerberg Biohub. L.L. is an investigator of the Howard Hughes Medical Institute. This work was supported by the National Institutes of Health (1K99-AG062746 to H.L., U24-CA210979 to D.R.M., U24-CA210986 to S.A.C., U01-CA214125 to S.A.C., R01-DK121409 to S.A.C. and A.Y.T., R01-CA186568 to A.Y.T., and R01-DC005982 to L.L.).

## AUTHOR CONTRIBUTIONS

J.L. and L.L. conceived this project. J.L., S.H., A.Y.T., and L.L. designed proteomic experiments. J.L., H.L., C.X., R.G., Q.X., T.L., D.J.L., and B.W. dissected fly brains. J.L. and S.H. processed proteomic samples with assistance from P.K. in BxxP synthesis. N.D.U., T.S., D.R.M., and S.A.C. performed post-enrichment sample processing, mass spectrometry, and initial data analysis. J.L. and S.H. analyzed proteomic data. J.L. and H.L. performed RNA sequencing on the platform provided by S.R.Q. J.L. designed and performed all the other experiments with input from L.L. and assistance from S.H., C.X., R.G., Q.X., D.J.L., C.N.M., and A.X. J.L., S.H., A.Y.T., and L.L. wrote the manuscript with input from all authors.

## STAR⋆METHODS

### KEY RESOURCES TABLE

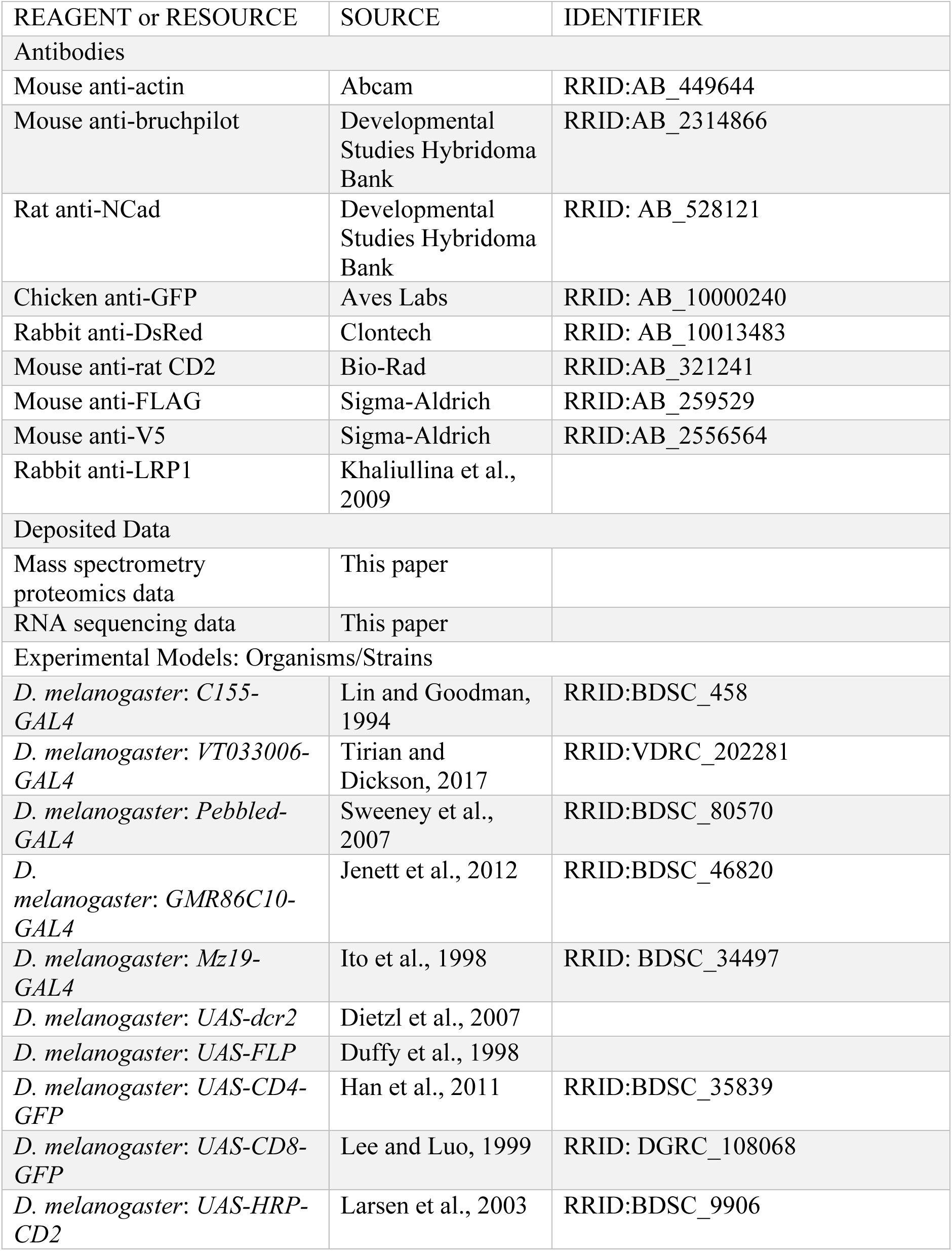
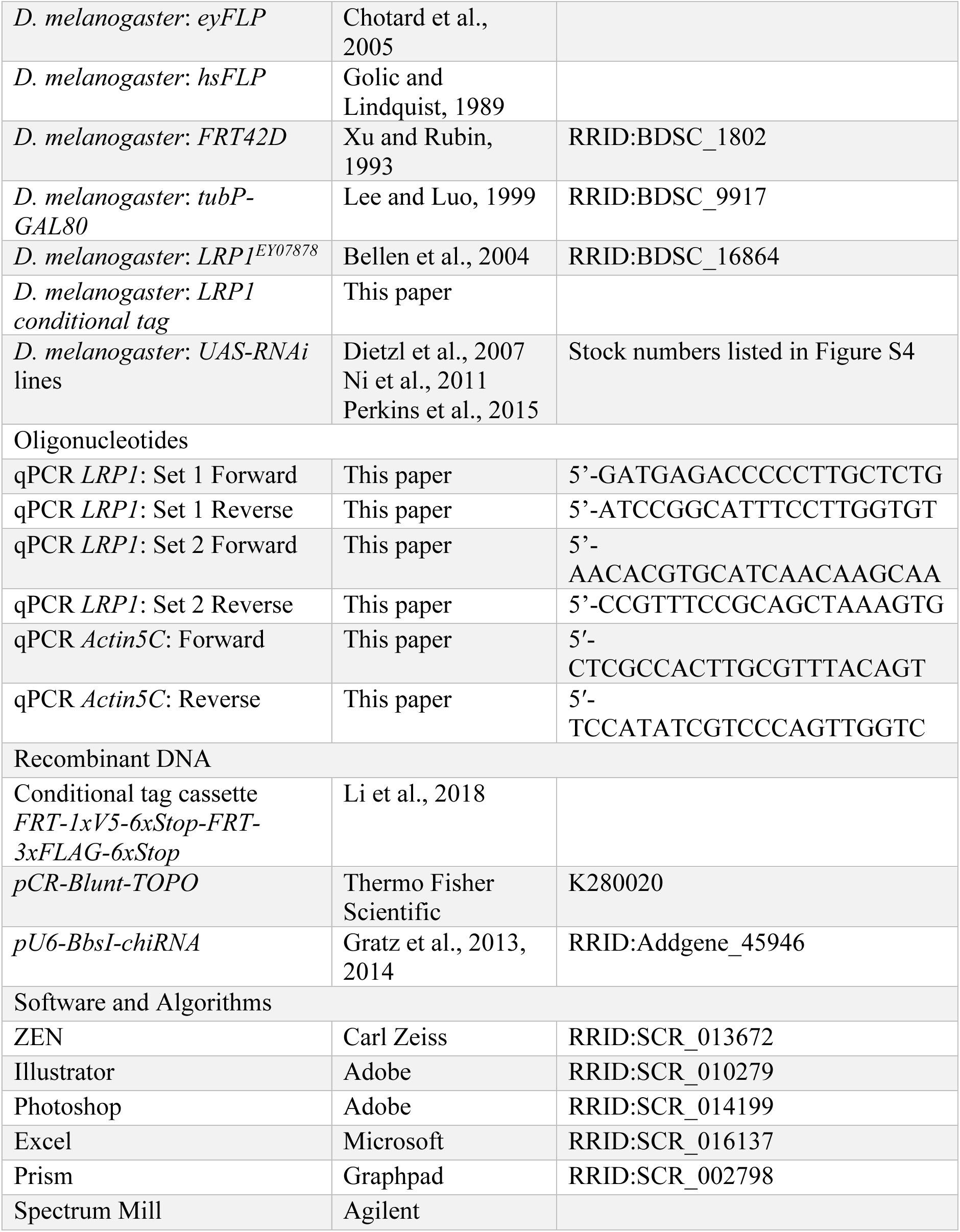

### CONTACT FOR REAGENT AND RESOURCE SHARING

Further information and requests for resources and reagents should be directed to and will be fulfilled by the Lead Contact, Liqun Luo.

### EXPERIMENTAL MODEL AND SUBJECT DETAILS

#### *Drosophila* stocks and genotypes

Flies were raised on standard cornmeal medium with a 12h/12h light cycle at 25°C, except for the RNAi experiments in which flies were raised at 29°C for enhanced transgenic expression. Complete genotypes of flies in each experiment are described in Table S1. The following lines were used: *C155-GAL4* (pan-neuronal *GAL4*) (Lin and Goodman, 1994), *VT033006-GAL4* (PN *GAL4*) (Tirian and Dickson, 2017), *Pebbled-GAL4* (ORN *GAL4*) (Sweeney et al., 2007), *GMR86C10-GAL4* (VM5d/v PN *GAL4*) (Jenett et al., 2012), *Mz19-GAL4* (DA1, VA1d, and DC3 PN *GAL4*) (Ito et al., 1998), *UAS-dcr2* (Dietzl et al., 2007), *UAS-FLP* (Duffy et al., 1998), *UAS-CD4-GFP* (Han et al., 2011), *UAS-CD8-GFP* (Lee and Luo, 1999), *UAS-HRP-CD2* (Larsen et al., 2003), *eyFLP* (ORN *FLP*) (Chotard et al., 2005), *hsFLP* (*heat shock protein* promoter-driven *FLP*) (Golic and Lindquist, 1989), *FRT42D* (Xu and Rubin, 1993), *tubP-GAL80* (*tubulin* promoter-driven *GAL80*) (Lee and Luo, 1999), *LRP1^EY07878^* (*LRP1* mutant; characterized in Figures S6A and S6B) (Bellen et al., 2004), and *LRP1 conditional tag* (Figure 5B; generated in this study, see below for details). The ventromedial (VM) genetic screen line (Figure 4B) was built previously (Xie et al., 2019). The RNAi lines were generated previously (Dietzl et al., 2007; Ni et al., 2011; Perkins et al., 2015) and acquired from the Bloomington *Drosophila* Stock Center, the Vienna *Drosophila* Resource Center, and the Japan National Institute of Genetics (stock numbers listed in Figure S4).

### METHOD DETAILS

#### BxxP synthesis

BxxP was synthesized as described previously (Loh et al., 2016). All chemicals were purchased from Sigma-Aldrich. Biotinamidohexanoyl-6-amino hexanoic acid N-hydroxysuccinimide ester (150 mg) and tyramine (36.2 mg) were dissolved in 2.7 mL of dimethylsulfoxide (DMSO). 276 μl (1.58 mmol, 6.0 equivalents) of DIPEA (N,N-diisopropylethylamine) was then added, and the reaction was incubated overnight at room temperature with stirring. 15 mL of H_2_O was added to quench the reaction before freezing and lyophilizing. 7 mL of cold methanol at −20°C was added drop-wise to the remaining brown-white solid mixture until the brown solid fully dissolved while the white material remained relatively insoluble. The solution was then chilled for 3 hours at −20°C. The white precipitate was separated using a fritted glass funnel and washed 4 times with 1 mL of ethyl acetate each time. After drying under vacuum, 117 mg of BxxP was obtained as a white solid. BxxP powder was subsequently dissolved in DMSO as 100 mM stock solution and stored in −20°C until use.

^1^H-NMR for BxxP (400 MHz, DMSO-d_6_): d = 9.17 (s, 1H), 7.89–7.62 (m, 3H), 6.97 (d, ^3^*J* = 8.4 Hz, 2H), 6.66 (d, ^3^*J* = 8.4 Hz, 2H), 6.43 (s, 1H), 6.37 (s, 1H), 4.36–4.26 (m, 1H), 4.19–4.08 (m, 1H), 3.23–2.89 (m, 6H), 2.81 (dd, ^2^*J* = 12.2 Hz, ^3^*J* = 5.2 Hz, 1H), 2.75–2.68 (m, 1H), 2.61–2.53 (m, 3H), 2.08–1.96 (m, 6H), 1.66–1.03 (m, 18H) ppm. LC/MS on an Agilent 6500 series Q-TOF: calculated for C_30_H_47_N_5_O_5_S [M+H]^+^: 590.33; found: 590.327.

#### Cell-surface biotinylation in fly brains

Dissection tools were thoroughly cleaned by Milli-Q ultrapure water (EMD Millipore) to remove detergent and other chemical contaminants. Brains were dissected in the Schneider’s medium (Thermo Fisher) pre-cooled on ice and transferred into 1.5 mL protein low-binding tubes (LoBind, Eppendorf) on ice, each containing 500 μL of the Schneider’s medium. Dissected brains were then briefly rinsed once with fresh Schneider’s medium to remove fat bodies and dissection debris. 100 μM BxxP was dissolved in fresh Schneider’s media by extensive vortex and sonication. Brains were then incubated with the BxxP-containing Schneider’s medium for 1 hour on ice, with occasional mixing via pipetting. 1 mM (0.003%) H_2_O_2_ (Thermo Fisher) was spiked into the medium for the 5-minute labeling reaction at room temperature. The reaction was then quenched immediately by five thorough washes with room temperature PBS (phosphate buffered saline; Thermo Fisher) containing 10 mM sodium ascorbate (Spectrum Chemicals), 5 mM Trolox (Sigma-Aldrich), and 10 mM sodium azide (Sigma-Aldrich). For biochemical characterization or proteomic sample collection, the quenching solution was drained, and the brains in minimal residual quenching solution were snap frozen in liquid nitrogen prior to storage at −80°C. For immunocytochemistry, the brains were immediately fixed and stained (see below for details).

#### Lysis of fly brains

Brains were processed in the original collection tube, to avoid loss during transferring. 40 μL of high-SDS RIPA buffer (50 mM Tris-HCl [pH 8.0], 150 mM NaCl, 1% sodium dodecyl sulfate (SDS), 0.5% sodium deoxycholate, 1% Triton X-100, 1x protease inhibitor cocktail (P8849), and 1 mM phenylmethylsulfonyl fluoride (PMSF); Sigma-Aldrich) was added to each tube of frozen brains. Disposable pestles driven by an electric motor (Thermo Fisher) were used to extensively grind the samples on ice. Brain lysates of the same experimental group were then merged into a single tube with a final volume of 300 μL of high-SDS RIPA buffer. Samples were then vortexed briefly, followed by two rounds of 10-second sonication (Branson 1800). To denature the post-synaptic density (Loh et al., 2016), samples were heated to 95°C for 5 minutes. 1.2 mL of SDS-free RIPA buffer (50 mM Tris-HCl [pH 8.0], 150 mM NaCl, 0.5% sodium deoxycholate, 1% Triton X-100, 1x protease inhibitor cocktail (P8849), and 1 mM PMSF) was added to each sample before 1-hour rotation at 4°C. Lysates were then transferred to 3.5 mL ultracentrifuge tubes (Beckman Coulter) containing 200 μL of normal RIPA buffer (50 mM Tris-HCl [pH 8.0], 150 mM NaCl, 0.2% SDS, 0.5% sodium deoxycholate, 1% Triton X-100, 1x protease inhibitor cocktail (P8849), and 1 mM PMSF) and centrifuged at 100,000 g for 30 minutes at 4°C. 1.5 mL of the supernatant was carefully collected for each sample and kept on ice.

#### Enrichment of biotinylated proteins

Streptavidin magnetic beads (Pierce) were used to enrich biotinylated proteins from brain lysates. For silver stain or Western blot, 20 μL was used for each sample from 50 dissected brains. For proteomic samples, 400 μL was used for each experimental group of 1000 dissected brains. Beads were first washed twice with normal RIPA buffer (50 mM Tris-HCl [pH 8.0], 150 mM NaCl, 0.2% SDS, 0.5% sodium deoxycholate, and 1% Triton X-100), and then incubated with the post-ultracentrifugation lysates on a 4°C rotator overnight. Beads were then sequentially washed twice with 1 mL of normal RIPA buffer, once with 1 mL of 1 M KCl, once with 1 mL of 0.1 M Na_2_CO_3_, once with 1 mL of 2 M urea in 10 mM Tris-HCl [pH 8.0], and twice with 1 mL of normal RIPA buffer. For silver stain or Western blot, biointylated proteins were eluted by heating the beads at 95°C for 10 minutes in 20 μL of 3x protein loading buffer (Bio-Rad) supplemented with 20 mM dithiothreitol (DTT) and 2mM biotin. For proteomic samples, on-bead trypsin digestion was performed after enrichment (see below for details). All chemicals were purchased from Sigma-Aldrich.

#### Silver stain and Western blot

4–12% Bis-Tris PAGE gels (Thermo Fisher) were used for protein electrophoresis following the manufacturer’s protocol. Silver stain kit (Pierce) was used for in-gel protein staining. For Western blot, proteins were transferred to nitrocellulose membranes (Thermo Fisher). All wash and incubation steps were performed on an orbital shaker at room temperature. After blocked with 3% bovine serum albumin (BSA) in TBST (Tris-buffered saline with 0.1% Tween 20; Thermo Fisher) for 1 hour, membranes were incubated with primary antibodies diluted in 3% BSA in TBST for 1 hour, followed by four rounds of 5-minute wash with TBST. Membranes were then incubated with horseradish peroxidase (HRP)-conjugated secondary antibodies diluted in 3% BSA in TBST for 1 hour, followed by four rounds of 5-minute wash with TBST. To blot biotinylated proteins, HRP-conjugated streptavidin was used instead of antibodies. Clarity Western ECL blotting substrate (Bio-Rad) and BioSpectrum imaging system (UVP) were used for chemiluminescence development and detection.

Primary antibodies used in this study include: mouse anti-actin (1:2000; ab8224, Abcam), mouse anti-bruchpilot (1:300; nc82, Developmental Studies Hybridoma Bank), and rat anti-NCad (1:300; DN-Ex#8, Developmental Studies Hybridoma Bank). HRP-conjugated secondary antibodies (Jackson ImmunoResearch or Thermo Fisher) were used at 1:3000. HRP-conjugated streptavidin (Thermo Fisher) was used at 0.3 μg/mL.

#### On-bead trypsin digestion of biotinylated proteins

To prepare proteomic samples for mass spectrometry analysis, proteins bound to streptavidin beads were washed twice with 200 μL of 50 mM Tris-HCl buffer [pH 7.5], followed by two washes with 2 M urea/50 mM Tris [pH 7.5] buffer. The final volume of 2 M urea/50 mM Tris [pH 7.5] buffer was removed, and beads were incubated with 80 μL of 2 M urea/50 mM Tris buffer containing 1 mM dithiothreitol (DTT) and 0.4 μg trypsin. Beads were incubated in the urea/trypsin buffer for 1 hour at 25°C while shaking at 1000 revolutions per minute (rpm). After 1 hour, the supernatant was removed and transferred to a fresh tube. The streptavidin beads were washed twice with 60 μL of 2 M urea/50 mM Tris [pH 7.5] buffer and the washes were combined with the on-bead digest supernatant. The eluate was reduced with 4 mM DTT for 30 minutes at 25°C with shaking at 1000 rpm. The samples were alkylated with 10 mM iodoacetamide and incubated for 45 minutes in the dark at 25°C while shaking at 1000 rpm. An additional 0.5 μg of trypsin was added to the sample and the digestion was completed overnight at 25°C with shaking at 700 rpm. After overnight digestion, the sample was acidified (pH < 3) by adding formic acid (FA) such that the sample contained ∼1% FA. Samples were desalted on C18 StageTips (3M). Briefly, C18 StageTips were conditioned with 100 μL of 100%MeOH, 100 μL of 50%MeCN/0.1% FA, and 2x with 100 μL of 0.1% FA. Acidified peptides were loaded onto the conditioned StageTips, which were subsequently washed 2x with 100 μL of 0.1%FA. Peptides were eluted from StageTips with 50 μL of 50%MeCN/0.1% FA and dried to completion.

#### TMT labeling and SCX StageTip fractionation of peptides

Desalted peptides were labeled with TMT (8-plex) reagents (Thermo Fisher) as directed by the manufacturer. Peptides were reconstituted in 100 μL of 50 mM HEPES. Each 0.8 mg vial of TMT reagent was reconstituted in 41 μL of anhydrous acetonitrile and added to the corresponding peptide sample for 1 hour at room temperature. Labeling of samples with TMT reagents was completed with the design described in Figure 2B. TMT labeling reactions were quenched with 8 μL of 5% hydroxylamine at room temperature for 15 minutes with shaking, evaporated to dryness in a vacuum concentrator, and desalted on C18 StageTips as described above.

For the TMT 8-plex cassette, 50% of the sample was fractionated by Strong Cation Exchange (SCX) StageTips while the other 50% of each sample was reserved for LC-MS analysis by a single shot, long gradient. One SCX StageTip was prepared per sample using 3 plugs of SCX material (3M) topped with 2 plugs of C18 material. StageTips were sequentially conditioned with 100 μL of MeOH, 100 μL of 80%MeCN/0.5% acetic acid, 100 μL of 0.5% acetic acid, 100 μL of 0.5% acetic acid/500mM NH_4_AcO/20% MeCN, followed by another 100 μL of 0.5% acetic acid. Dried sample was re-suspended in 250 μL of 0.5% acetic acid, loaded onto the StageTips, and washed 2x with 100 μL of 0.5% acetic acid. Sample was trans-eluted from C18 material onto the SCX with 100 μL of 80%MeCN/0.5% acetic acid, and consecutively eluted using 3 buffers with increasing pH. The first elution was with pH 5.15 (50mM NH_4_AcO/20% MeCN), followed by pH 8.25 (50mM NH_4_HCO_3_/20% MeCN), and finally with pH 10.3 (0.1% NH_4_OH, 20% MeCN). Three eluted fractions were re-suspended in 200 μL of 0.5% acetic acid, to reduce the MeCN concentration, and subsequently desalted on C18 StageTips as described above. Desalted peptides were dried to completion.

#### Liquid chromatography and mass spectrometry

Desalted, TMT-labeled peptides were resuspended in 9 μL of 3% MeCN, 0.1% FA and analyzed by online nanoflow liquid chromatography tandem mass spectrometry (LC-MS/MS) using a Fusion Lumos mass spectrometer (Thermo Fisher) coupled on-line to a Proxeon Easy-nLC 1000 (Thermo Fisher). Four microliters of each sample was loaded at 500 nl/min onto a microcapillary column (360 μm outer diameter × 75 μm inner diameter) containing an integrated electrospray emitter tip (10 μm), packed to approximately 24 cm with ReproSil-Pur C18-AQ 1.9 μm beads (Dr. Maisch GmbH) and heated to 50 °C. The HPLC solvent A was 3% MeCN, 0.1% FA, and the solvent B was 90% MeCN, 0.1% FA. Peptides were eluted into the mass spectrometer at a flow rate of 200 nl/min. The SCX fractions were run with 110-minute method, which used the following gradient profile: (min:%B) 0:2; 1:6; 85:30; 94:60; 95:90; 100:90; 101:50; 110:50 (the last two steps at 500 nL/min flow rate). Non-fractionated samples were analyzed using a 260 min LC-MS/MS method with the following gradient profile: (min:%B) 0:2; 1:6; 235:30; 244:60; 245:90; 250:90; 251:50; 260:50 (the last two steps at 500 nL/min flow rate). The Fusion Lumos was operated in the data-dependent mode acquiring HCD MS/MS scans (r =50,000) after each MS1 scan (r = 60,000) on the top 12 most abundant ions using an MS1 target of 3 × 106 and an MS2 target of 5 × 104. The maximum ion time utilized for MS/MS scans was 120 ms; the HCD normalized collision energy was set to 34; the dynamic exclusion time was set to 20 s, and the peptide match and isotope exclusion functions were enabled. Charge exclusion was enabled for charge states that were unassigned, 1 and >7.

#### Mass spectrometry data processing

Collected data were analyzed using the Spectrum Mill software package v6.1 pre-release (Agilent Technologies). Nearby MS scans with the similar precursor m/z were merged if they were within ± 60 s retention time and ±1.4 m/z tolerance. MS/MS spectra were excluded from searching if they failed the quality filter by not having a sequence tag length 0 or did not have a precursor MH+ in the range of 750 – 4000. All extracted spectra were searched against a UniProt database containing *Drosophila melanogaster* reference proteome sequences. Search parameters included: ESIQEXACTIVE-HCD-v2 scoring parent and fragment mass tolerance of 20 ppm, 30% minimum matched peak intensity, trypsin allow P enzyme specificity with up to four missed cleavages, and calculate reversed database scores enabled. Fixed modifications were carbamidomethylation at cysteine. TMT labeling was required at lysine, but peptide N termini were allowed to be either labeled or unlabeled. Allowed variable modifications were protein N-terminal acetylation and oxidized methionine. Individual spectra were automatically assigned a confidence score using the Spectrum Mill auto-validation module. Score at the peptide mode was based on target-decoy false discovery rate (FDR) of 1%. Protein polishing auto-validation was then applied using an auto thresholding strategy. Relative abundances of proteins were determined using TMT reporter ion intensity ratios from each MS/MS spectrum and the median ratio was calculated from all MS/MS spectra contributing to a protein subgroup. Proteins identified by 2 or more distinct peptides and ratio counts were considered for the dataset.

#### Proteomic data analysis

To determine the cutoff in each biological replicate, we adopted the ratiometric analysis as previously described (Hung et al., 2014). Detected proteins were classified according to the annotation of subcellular localization in the UniProt database (retrieved in March 2018). Proteins with the plasma membrane annotation were classified as true-positives (TPs). Proteins with either nucleus, mitochondrion, or cytoplasm annotation but without the membrane annotation were classified as false-positives (FPs). Of the total 2020 detected proteins, 335 were TPs and 628 were FPs. For each replicate, the proteins were first ranked in a descending order according to the TMT ratio (127C/130C, 129N/131, 129C/127N, or 126/128C). For each protein on the ranked list, the accumulated true-positive count and false-positive count above its TMT ratio were calculated. A receiver operating characteristic (ROC) curve was plotted accordingly for each replicate (Figure 2D). The cutoff was set where *true-positive rate – false-positive rate* (TPR – FPR) maximized (Figure S2A): 127C/130C: 0.038; 129N/131: 0.197; 129C/127N: 0.153; and 126/128C: 0.121. Post-cutoff proteomic lists of the two biological replicates for each time point were intersected to obtain the final proteome. We also performed the cutoff analysis with a different TMT pairing regime (127C/131, 129N/130C, 129C/128C, and 126/127N) and obtained almost identical proteomes (data not shown).

For gene ontology analysis (Figures 2G and 3A), we uploaded the final proteomes to the STRING database search portal and plotted the top five retrieved terms on cellular compartment or biological process with the lowest false discovery rates. For the protein network (Figure 3E), proteins with known function in neural development or synaptic transmission were clustered by their reported protein-protein interactions and corresponding confidence scores (STRING) using a Markov clustering algorithm (inflation value set at 2.5) and plotted in Cytoscape (v3.7.1). FlyBase and NCBI PubMed were searched to determine if each protein in the final proteomes had reported function in neural development or synaptic transmission.

#### Quantitative comparison of developing and mature proteomes

For the volcano plot (Figures 3D and 4A) comparing differentially enriched proteins in developing versus mature samples, a linear model was fit to account for the variance across replicates for each stage and normalize data by the appropriate negative control samples. A protein summary was first generated where each TMT condition was calculated as a ratio to the median intensity of all the channels, enabling all channels to have the same denominator. Then, for each protein, a linear model was used to calculate the following ratio and the corresponding *p* value:

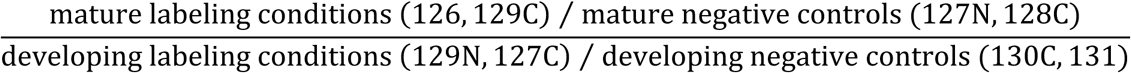

Using log_2_ transformed TMT ratios, the linear model is as follows:

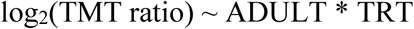

where ADULT and TRT are indicator variables representing maturity (ADULT=1 for mature, 0 for developing) and labeling condition (TRT=1 for labeled, 0 for negative control) respectively. The above linear model with interaction terms expands to:

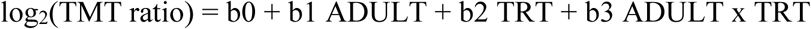

Coefficient b3 represents the required (log-transformed) ratio between mature and developing conditions taking into account the appropriate negative controls and replicates. A moderated *t*-test was used to test the null hypothesis of b3=0 and calculate a nominal *p*-value for each protein. These nominal *p*-values were then corrected for multiple testing using the Benjamini-Hochberg FDR (BH-FDR) method (Benjamini and Hochberg, 1995). The linear model along with the associated moderated *t*-test and BH-FDR correction were implemented using the limma library (Ritchie et al., 2015) in R.

#### RNA sequencing and data processing

Fly brains with olfactory PNs labelled by *VT033006-GAL4>CD8-GFP* were dissected in the Schneider’s medium (Thermo Fisher). Single-cell suspensions were then prepared as previously described (Li et al., 2017). Briefly, the dissected brains were dissociated by papain (Worthington) and liberase TM (Roche), along with extensive pipetting and needle passing. Two thousand GFP+ cells were collected for each sample using fluorescence activated cell sorting (FACS) on a Sony SH800 cell sorter system (Sony Biotechnology). Full-length poly(A)-tailed RNA was reverse-transcribed and amplified by PCR, following a modified SMART-seq2 protocol (Li et al., 2017; Picelli et al., 2014). Sequencing libraries were prepared from amplified cDNA using tagmentation (Nextera XT, Illumina). Sequencing was performed using the Nextseq 500 platform (Illumina) with paired-end 75 bp reads.

Reads were aligned to the *Drosophila melanogaster* genome (r6.10) using STAR (2.4.2a) (Dobin et al., 2013) with the ENCODE standard options, except “–outFilterScoreMinOverLread 0.4–outFilterMatchNminOverLread 0.4–outFilterMismatchNmax 999–outFilterMismatchNoverLmax 0.04”. Uniquely mapped reads that overlap with genes were counted using HTSeq-count (0.7.1) (Anders et al., 2015) with default settings except “–m intersection–strict”. To normalize for comparison among samples, we rescaled gene counts to counts per million (CPM). All analyses were performed after converting gene counts to logarithmic space via the transformation log_2_(CPM+1).

#### Generation of *LRP1* conditional tag

The *FRT-1xV5-6xStop-FRT-3xFLAG-6xStop* cassette was amplified from the vector for generating *PlexB conditional tag* (Li et al., 2018). A *loxP*-flanked *miniWhite* cassette was inserted in front of the second *FRT* site, to build the *FRT-1xV5-6xStop-loxP-miniWhite-loxP-FRT-3xFLAG-6xStop* cassette (hereafter, conditional tag cassette). *Drosophila* genomic DNA was extracted from *w^1118^* adult flies by DNeasy blood and tissue kit (Qiagen). Genomic sequence flanking the endogenous stop codon of *LRP1* was amplified by Q5 hot-start high-fidelity DNA polymerase (New England Biolabs) and inserted into a *pCR-Blunt-TOPO* vector by Blunt TOPO PCR cloning kit (Thermo Fisher). The conditional tag cassette was inserted into the *pCR-LRP1 genomic sequence* plasmid to replace the endogenous stop codon by NEBuilder HiFi DNA assembly master mix (New England Biolabs), thus building the CRISPR HDR (homology directed repair) vector – *pCR-LRP1 coding sequence-conditional tag cassette-LRP1 3’UTR*. CRISPR guide RNA (gRNA) targeting a locus near *LRP1* stop codon was designed by the flyCRISPR Target Finder tool and cloned into a *pU6-BbsI-chiRNA* vector (Gratz et al., 2013, 2014). The HDR and *pU6-gRNA* plasmids were transformed into NEB stable competent *E. coli* (New England Biolabs), extracted by QIAprep spin miniprep kit (Qiagen), and verified by full-length sequencing (Elim Biopharmaceuticals). The verified HDR and *pU6-gRNA* plasmids were co-injected into *vasa-Cas9* (Port et al., 2014) fly embryos. All *white+* progenies were individually balanced by *CyO*. The *loxP*-flanked *miniWhite* cassette was then removed by crossing each line to a *heat shock protein* promoter-driven Cre (*hs-Cre*). After multiple sessions of 2-hour heat shock at 37°C, the *white^−^* progenies were individually balanced by *CyO* and verified by sequencing the HDR-covered genomic region.

#### Quantitative real-time PCR

Total RNA of *w^1118^* and *LRP1* mutant (*LRP1^EY07878^*) flies was extracted from 2nd-instar larvae by PureLink TRIzol RNA mini kit (Thermo Fisher). Two sets of primers targeting non-overlapping regions of *LRP1* cDNA (Set 1: 5’-GATGAGACCCCCTTGCTCTG and 5’-ATCCGGCATTTCCTTGGTGT; Set 2: 5’-AACACGTGCATCAACAAGCAA and 5’-CCGTTTCCGCAGCTAAAGTG) were designed by NCBI Primer-BLAST. Quantitative real-time PCR was performed with the iTaq universal SYBR Green one-step kit (Bio-Rad) on a CFX96 real-time PCR detection system (Bio-Rad). The *LRP1* mRNA level was normalized to the *Act5C* mRNA level in each sample (*Act5C* primers: 5′-CTCGCCACTTGCGTTTACAGT and 5′-TCCATATCGTCCCAGTTGGTC).

#### MARCM-based mosaic analysis

MARCM analyses were performed as previously described (Lee and Luo, 1999; Wu and Luo, 2006a; Yu et al., 2010). Complete fly genotypes of MARCM experiments are described in Table S1. In *hsFLP*- and *GMR86C10-GAL4*-based MARCM of VM5d/v PNs, larvae and early-stage pupae (72 to 120 hours after hatching) were heat shocked for 1.5 hours at 37°C. In *hsFLP*- and *Mz19-GAL4*-based MARCM of DA1, VA1d, and DC3 PNs, larvae (24 to 48 hours after hatching) were heat shocked for 1 hour at 37°C. Brains of adult flies were dissected, immunostained, and imaged as described below.

#### Immunocytochemistry

Dissection and immunostaining (except the staining of LRP1 conditional tag, see below for details) of fly brains were performed according to previously described methods (Wu and Luo, 2006b). Briefly, the brains were dissected in PBS (phosphate buffered saline; Thermo Fisher) and then fixed in 4% paraformaldehyde (Electron Microscopy Sciences) in PBS with 0.015% Triton X-100 (Sigma-Aldrich) for 20 minutes on a nutator at room temperature. Fixed brains were washed with PBST (0.3% Triton X-100 in PBS) four times, each time nutating for 20 minutes. The brains were then blocked in 5% normal donkey serum (Jackson ImmunoResearch) in PBST for 1 hour at room temperature or overnight at 4°C on a nutator. Primary antibodies were diluted in the blocking solution and incubated with brains for 36-48 hours on a 4°C nutator. After washed with PBST four times, each time nutating for 20 minutes, brains were incubated with secondary antibodies diluted in the blocking solution and nutated in the dark for 36-48 hours at 4°C. Brains were then washed again with PBST four times, each time nutating for 20 minutes. Immunostained brains were mounted with SlowFade antifade reagent (Thermo Fisher) and stored at 4°C before imaging.

For the staining of LRP1 conditional tag, the routine protocol described above failed to detect FLAG or V5 signal from the background, likely due to the low expression of endogenous LRP1 proteins *in vivo*. Alexa 488 Tyramide SuperBoost kit (Thermo Fisher) was used to amplify the immunostaining signal by following the manufacture’s protocol. Briefly, the brains were dissected, fixed, and washed as described above. After rinsed with PBS twice, the brains were incubated with 3% hydrogen peroxide for 1 hour at room temperature to quench the activity of endogenous peroxidases, and then washed with PBS three times. After blocked in 10% goat serum for 1 hour at room temperature, the brains were nutated in primary antibodies diluted in 10% goat serum for 36–48 hours at 4°C. After washed with PBST four times, each time nutating for 20 minutes, the brains were incubated with the poly-HRP-conjugated secondary antibody provided in the kit and nutated for 36–48 hours at 4°C. Then, the brains were washed with PBST four times, each time nutating for 20 minutes, followed by two rounds of fast rinsing in PBS. The tyramide working solution and the quenching buffer were made freshly according to the kit’s recipe. The brains were incubated with the tyramide solution for 6 minutes at room temperature and immediately washed with the quenching buffer three times, followed by four rounds of thorough washing with PBST. Stained brains were mounted with SlowFade antifade reagent (Thermo Fisher) and stored at 4°C before imaging.

Primary antibodies used in immunostaining include: rat anti-NCad (1:40; DN-Ex#8, Developmental Studies Hybridoma Bank), chicken anti-GFP (1:1000; GFP-1020, Aves Labs), rabbit anti-DsRed (1:200; 632496, Clontech), mouse anti-rat CD2 (1:200; OX-34, Bio-Rad), mouse anti-FLAG (1:100; M2, Sigma-Aldrich), mouse anti-V5 (1:100; R960-25, Thermo Fisher), and rabbit anti-LRP1 (1:200; gift of Suzanne Eaton) (Khaliullina et al., 2009). Donkey secondary antibodies conjugated to Alexa Fluor 405/488/568/647 (Jackson ImmunoResearch or Thermo Fisher) were used at 1:250. Neutravidin (Thermo Fisher) pre-conjugated with Alexa Fluor 647 was used to detect biotin.

#### Image acquisition and processing

Images were acquired by a Zeiss LSM 780 laser-scanning confocal microscope (Carl Zeiss), with a 40x/1.4 Plan-Apochromat oil objective (Carl Zeiss). Confocal z-stacks were obtained by 1-μm intervals at the resolution of 512×512. Images were exported as maximum projections or single confocal sections by ZEN (Carl Zeiss) in the format of TIFF. Photoshop (Adobe) was used for image rotation and cropping. BioRender was used to make diagrams. Illustrator (Adobe) was used to assemble figures.

#### Statistics and data analysis

No statistical methods were used to determine sample sizes, but our sample sizes were similar to those generally employed in the field. Antennal lobes damaged in dissection were excluded from analysis; otherwise, all samples were included. Data collection and analysis were not performed blind to the conditions of the experiments. Excel (Microsoft) and Prism (GraphPad) were used for data analysis and plotting. Statistical methods used were described in the figure legend of each relevant panel.

## DATA AND SOFTWARE AVAILABILITY

Original proteomic data prior to analyses is provided in Table S2. Mass spectrometry and sequencing data will be uploaded to public data depositories before publication.

---

**Figure S1.**
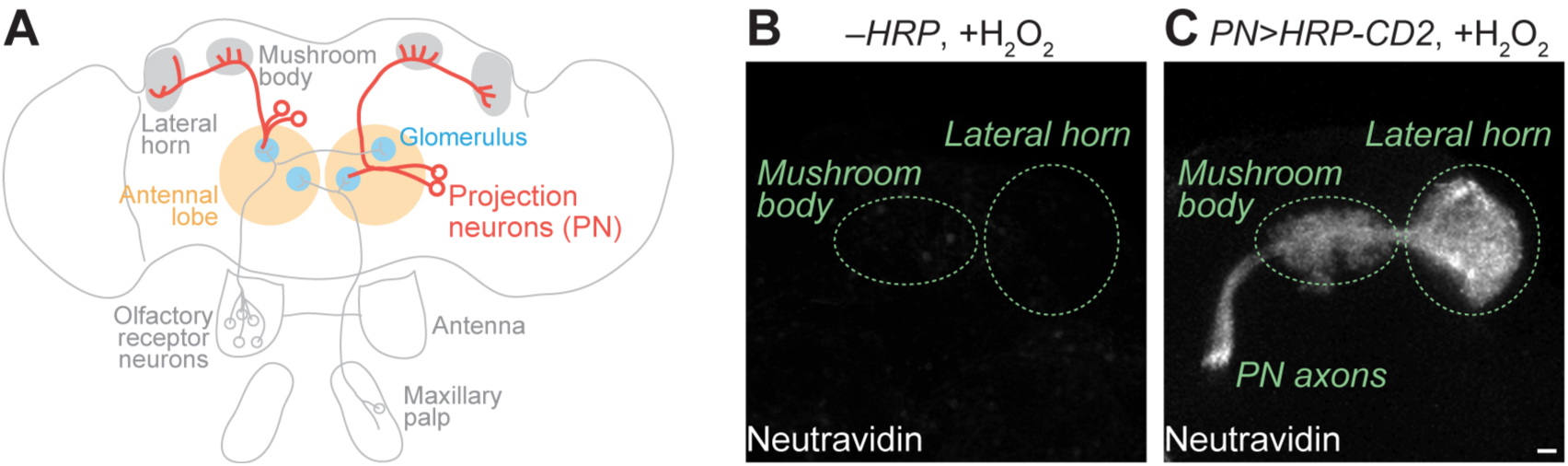
Cell-surface biotinylation of olfactory PNs in intact brains. Related to Figure 1. (**A**) Scheme of the fly olfactory circuit. (**B** and **C**) Neutravidin staining of PN axon-targeted brain regions after the cell-surface biotinylation reaction. (B) HRP was not expressed by omitting the *GAL4* driver. (C) *PN-GAL4* drove the expression of cell surface-targeted HRP (HRP-CD2). Scale bar, 10 μm.

**Figure S2.**
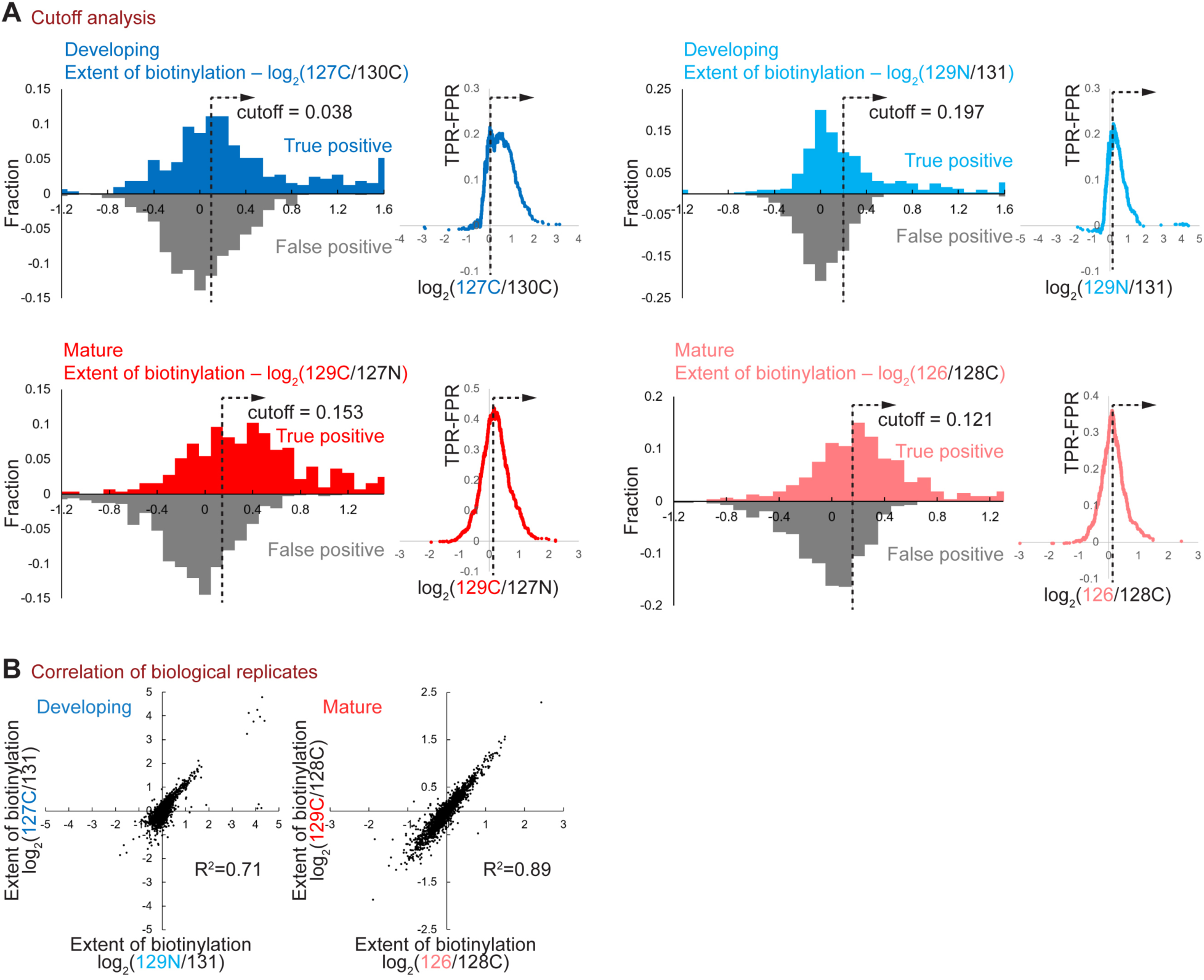
Analysis of PN surface proteomes. Related to Figure 2. (**A**) Determination of the TMT ratio cutoff in each biological replicate. Cutoffs were set where *true-positive rate – false-positive rate* (TPR – FPR) maximized. True-positive denotes plasma membrane proteins curated by the UniProt database. False-positive includes nuclear, mitochondrial, and cytosolic proteins without membrane annotation by UniProt. (**B**) Correlation of biological replicates using negative controls different from Figure 2F.

**Figure S3.**
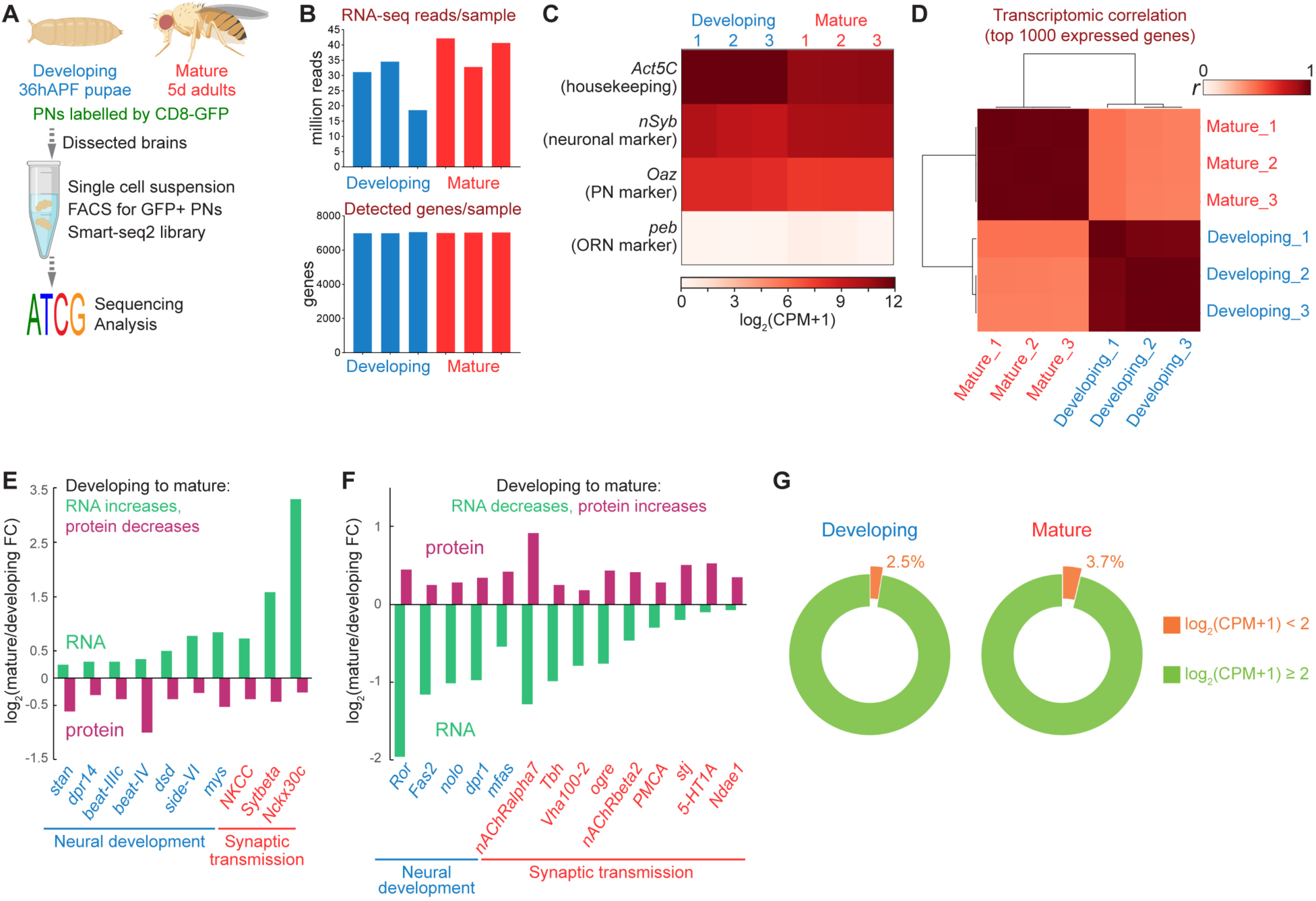
RNA vs. protein dynamics of PN surface molecules. Related to Figure 3. (**A**) Workflow of the bulk PN RNA sequencing. (**B**) The read number and detected gene number in each of the three biological replicates for both stages. (**C**) Expression levels of marker genes in RNA sequencing. CPM, counts per million. (**D**) Transcriptomic correlation of biological replicates, calculated by the top 1000 expressed genes. (**E** and **F**) Many functionally critical proteins in neural development or synaptic transmission exhibited inverse dynamics in RNA sequencing and cell-surface protein profiling. In the developing-to-mature transition, RNA increased (log_2_FC > 0.1) but protein decreased (log_2_FC < – 0.1) (E) or RNA decreased (log_2_FC < –0.1) but protein increased (log_2_FC > 0.1) (F). FC, fold change. (**G**) A small fraction (orange) of proteins captured on the PN surface showed very low read counts in PN RNA-sequencing. CPM, counts per million.

**Figure S4.**
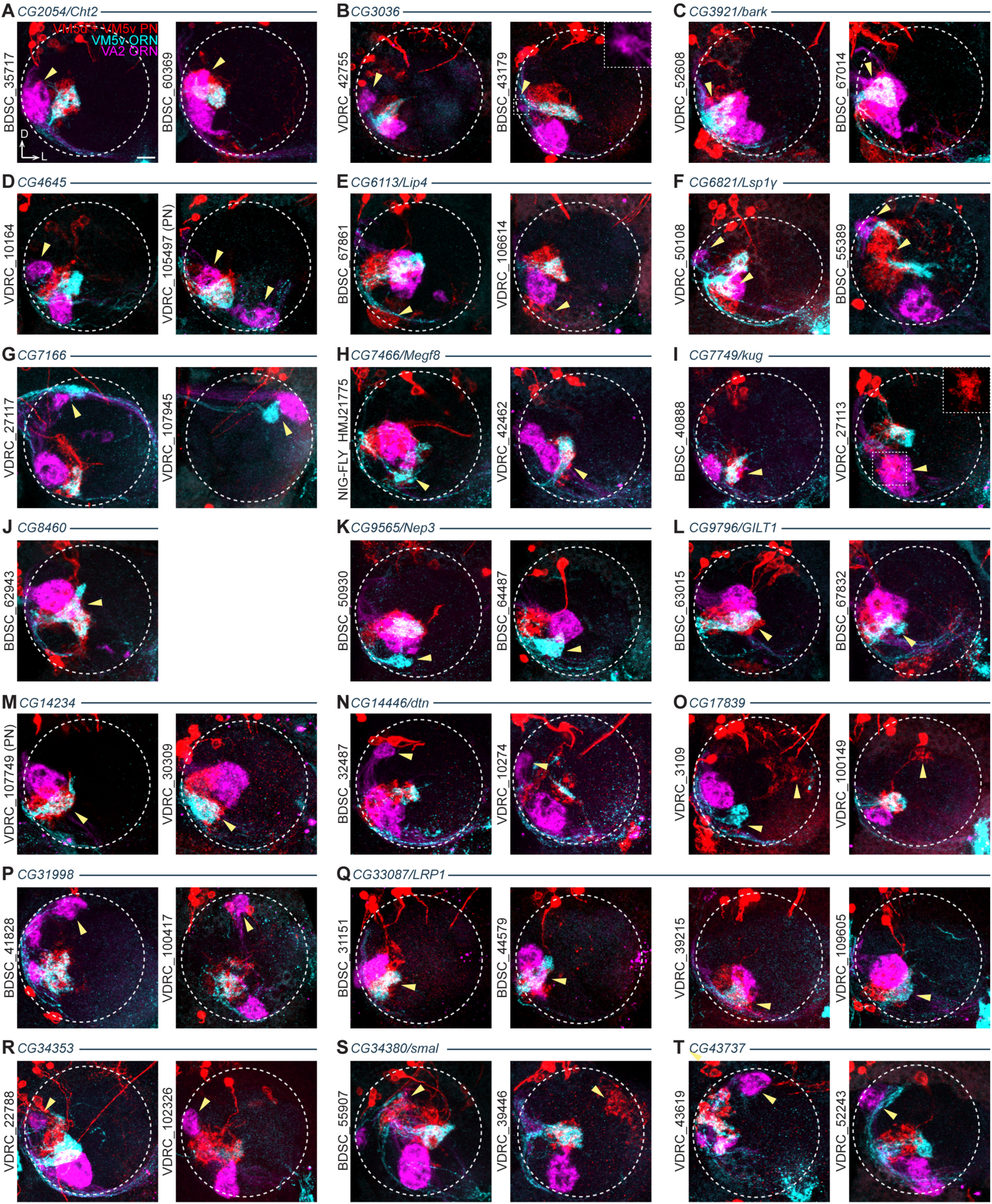
RNAi phenotypes of the genetic screen in VM PNs and ORNs. Related to Figure 4 and Table 2. Design of the genetic screen is described in Figure 4B. The primary (most penetrant) phenotypes are described below. Phenotypic penetrances are listed in Table 2. Arrowhead, mistargeted PN dendrites or ORN axons. Dashed circle, antennal lobe. (**A**) *CG2054/Cht2*, medial mistargeting of VA2 ORN axons. (**B**) *CG3036*, posterior-medial mistargeting of VA2 ORN axons. (**C**) *CG3921/bark*, local mistargeting of VM5d/v PN dendrites and VM5v/VA2 ORN axons. (**D**) *CG4645*, medial mistargeting of VA2 ORN axons. VDRC_105497 was lethal when driven by the pan-neuronal *C155-GAL4*. *PN-GAL4* was used instead. (**E**) *CG6113/Lip4*, ventral mistargeting of VM5d/v PN dendrites. (**F**) *CG6821/Lsp1γ*, global disruption of the antennal lobe structure and randomized PN dendrite/ORN axon targeting. (**G**) *CG7166*, ectopic dorsolateral trajectory and dorsal mistargeting of VM5v/VA2 ORN axons. (**H**) *CG7466/Megf8*, ventral mistargeting of VM5v ORN axons. (**I**) *CG7749/kug*, ventral mistargeting of VM5d/v PN dendrites. (**J**) *CG8460*, dorsal mistargeting of VA2 ORN axons. Only one RNAi line was available. (**K**) *CG9565/Nep3*, ventral mistargeting of VM5v ORN axons. (**L**) *CG9796/GILT1*, locally dorsal shifting of VA2 ORN axons, as well as aberrant extension of VM5d/v PN dendrites and VM5v ORN axons. (**M**) *CG14234*, locally dorsal shifting of VA2 ORN axons, as well as locally ventral shifting of VM5d/v PN dendrites and VM5v ORN axons. VDRC_107749 was lethal when driven by the pan-neuronal *C155-GAL4*. *PN-GAL4* was used instead. (**N**) *CG14446/dtn*, dorsal mistargeting of VA2 ORN axons. (**O**) *CG17839*, lateral mistargeting of VM5d/v PN dendrites. (**P**) *CG31998*, dorsal mistargeting of VA2 ORN axons. (**Q**) *CG33087/LRP1*, local mistargeting of VM5d/v PN dendrites and VM5v/VA2 ORN axons. (**R**) *CG34353*, posterior-medial mistargeting of VA2 ORN axons. (**S**) *CG34380/small*, random long-range mistargeting of VM5d/v PN dendrites. (**T**) *CG43737*, dorsal mistargeting of VM5v and VA2 ORN axons, as well as random mistargeting of VM5d/v PN dendrites. Scale bar, 10 μm. D, dorsal; L, lateral.

**Figure S5.**
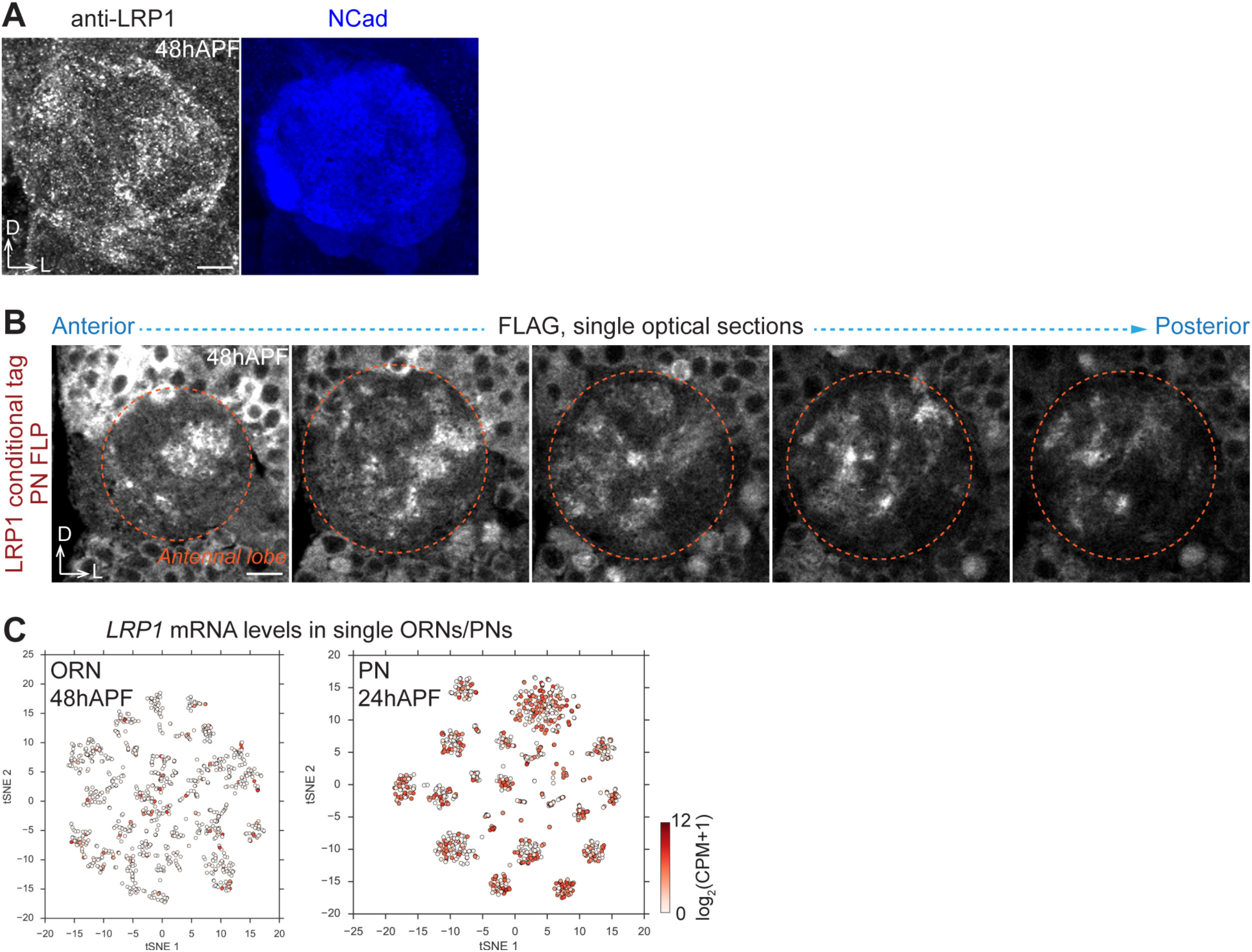
Expression pattern of LRP1. Related to Figure 5. (**A**) LRP1 antibody staining of the antennal lobe at 48 hours after puparium formation (48hAPF). NCad, neuropil. (**B**) Single optical sections of FLAG staining of LRP1 conditional tag with PN-specific FLP at 48hAPF. Orange circle, antennal lobe. (**C**) *LRP1* mRNA levels in single ORNs (48hAPF) (Li et al., 2019) and PNs (24hAPF) (Li et al., 2017). CPM, counts per million. Scale bar, 10 μm. D, dorsal; L, lateral.

**Figure S6.**
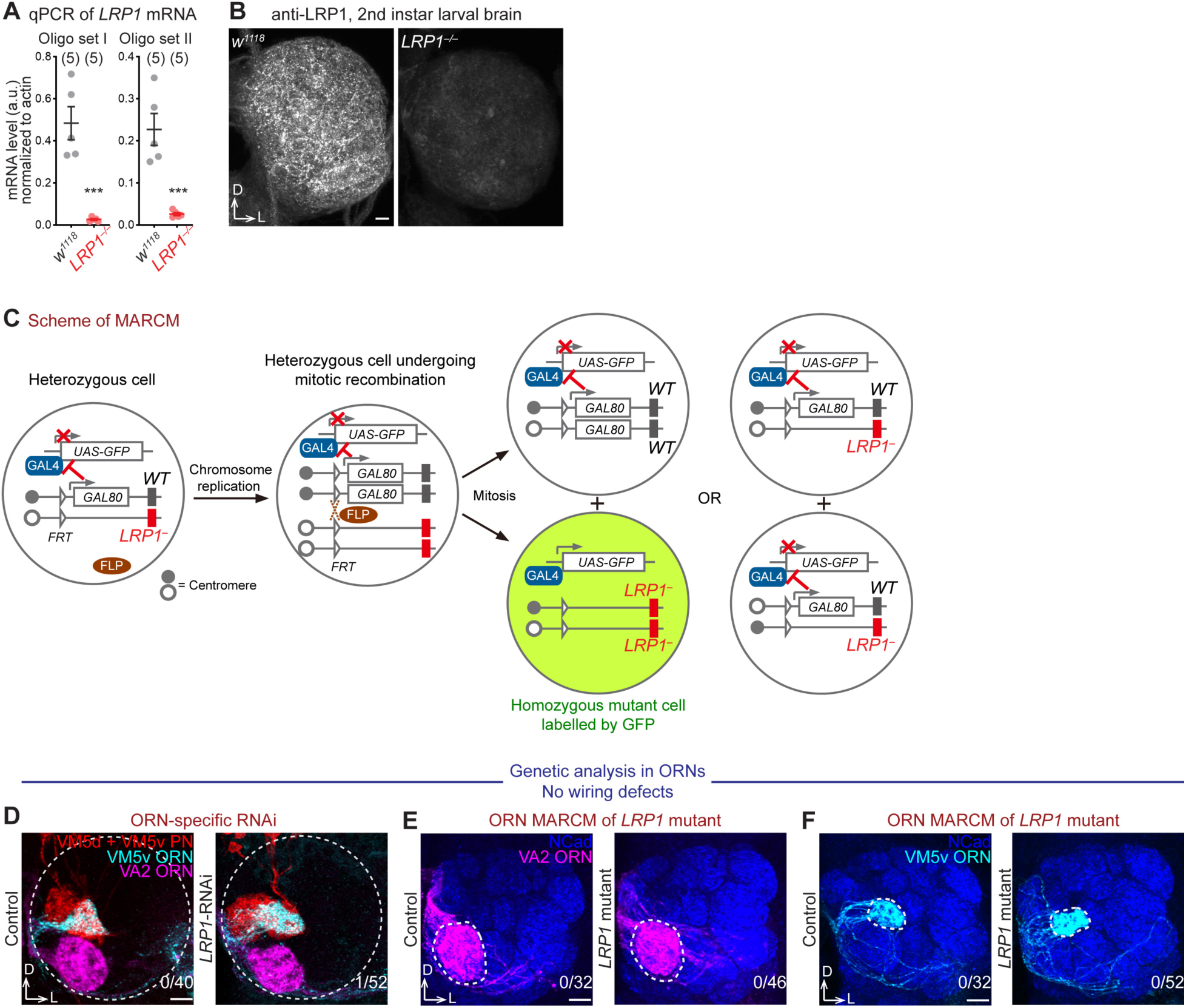
MARCM-based mosaic analysis of *LRP1* null mutant. Related to Figure 5. (**A**) Quantification of *LRP1* mRNA levels in *wild-type* (*w^1118^*) and *LRP1* homozygous mutant (*LRP1^EY07878/EY07878^*, noted as *LRP1^™/™^*) fly larvae. Error bars denote standard errors. An unpaired, two-tailed *t*-test was used to determine the statistical significance. ***, *p*<0.001. (**B**) LRP1 antibody staining of *wild-type* (*w^1118^*) and *LRP1* homozygous mutant (*LRP1^™/™^*) fly larval brains. (**C**) Scheme of MARCM (mosaic analysis with a repressible cell marker). GAL80 suppresses the GFP expression driven by the GAL4*/UAS* binary system. The mutant (*LRP1^−^*) allele is on the same chromosome arm in *trans* to the *GAL80* transgene. After FLP-mediated mitotic recombination, only the homozygous mutant cell loses *GAL80* and is thus labelled by GFP. (**D**) ORN-specific *LRP1*-RNAi knockdown did not cause any wiring defects in VM PN dendrites or ORN axons. (**E** and **F**) Mosaic loss of *LRP1* only in ORNs using *eyFLP*-based MARCM did not show any wiring defects in VA2 (E) or VM5v (F) ORN axons. Scale bar, 10 μm. D, dorsal; L, lateral. The number of antennal lobes with mistargeting over the total number of antennal lobes examined is noted at the bottom-right corner of each panel.

**Table S1.**
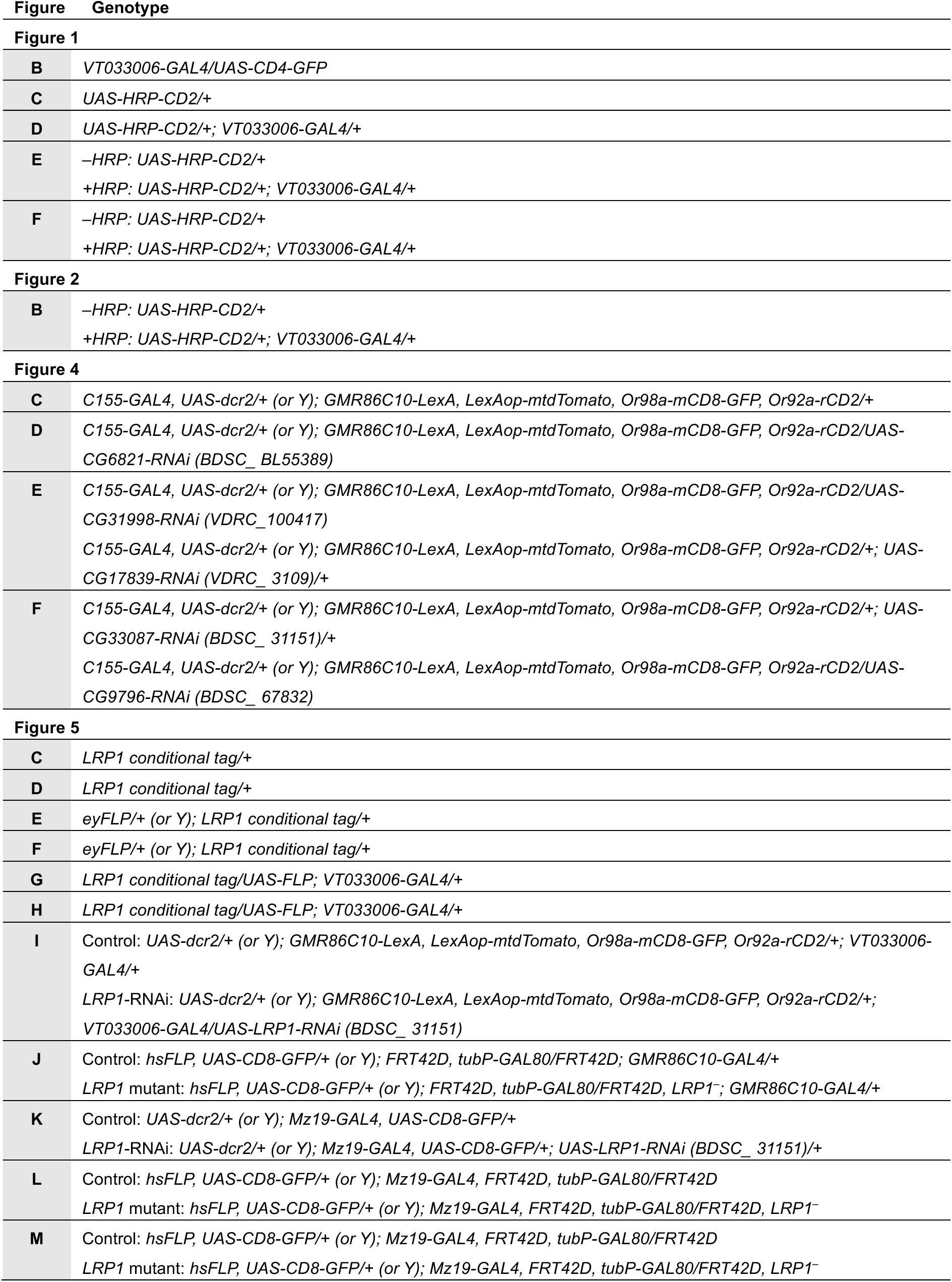
Genotypes of flies in each experiment.

**Table S2.**
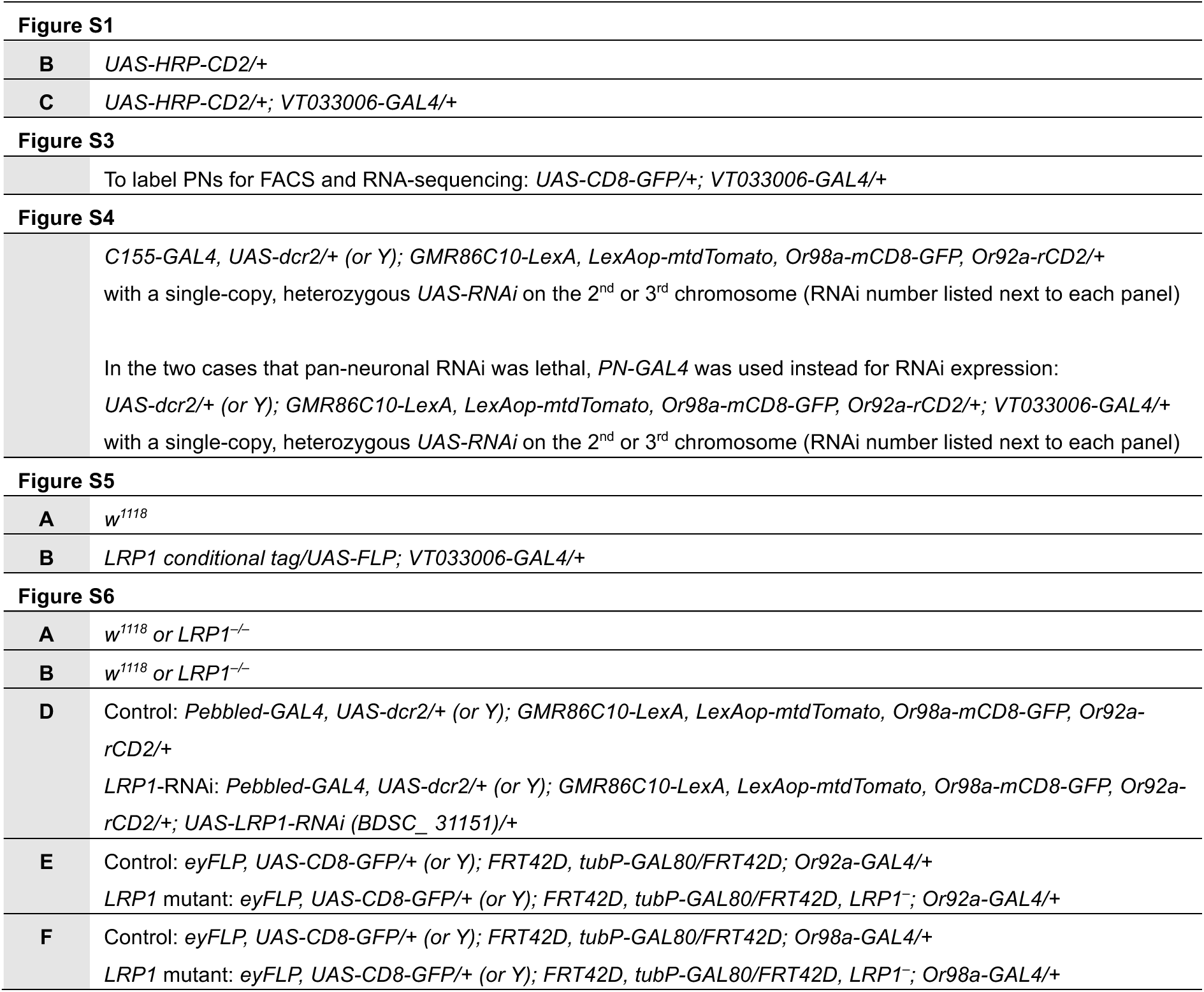
Spreadsheet of original proteomic data prior to analyses.

## REFERENCES

Aebersold, R., and Mann, M. (2016). Mass-spectrometric exploration of proteome structure and function. Nature 537, 347–355.

Almagro Armenteros, J.J., Sønderby, C.K., Sønderby, S.K., Nielsen, H., and Winther, O. (2017). DeepLoc: prediction of protein subcellular localization using deep learning. Bioinformatics 33, 3387–3395.

Alvarez-Castelao, B., Schanzenbächer, C.T., Hanus, C., Glock, C., Tom Dieck, S., Dörrbaum, A.R., Bartnik, I., Nassim-Assir, B., Ciirdaeva, E., Mueller, A., et al. (2017). Cell-type-specific metabolic labeling of nascent proteomes in vivo. Nat. Biotechnol. 35, 1196–1201.

Anders, S., Pyl, P.T., and Huber, W. (2015). HTSeq — a Python framework to work with high-throughput sequencing data. Bioinformatics 31, 166–169.

Barish, S., Nuss, S., Strunilin, I., Bao, S., Mukherjee, S., Jones, C.D., and Volkan, P.C. (2018). Combinations of DIPs and Dprs control organization of olfactory receptor neuron terminals in Drosophila. PLoS Genet. 14, e1007560.

Bellen, H.J., Levis, R.W., Liao, G., He, Y., Carlson, J.W., Tsang, G., Evans-Holm, M., Hiesinger, P.R., Schulze, K.L., Rubin, G.M., et al. (2004). The BDGP gene disruption project: single transposon insertions associated with 40% of Drosophila genes. Genetics 167, 761–781.

Benjamini, Y., and Hochberg, Y. (1995). Controlling the false discovery rate: A practical and powerful approach to multiple testing. Journal of the Royal Statistical Society: Series B (Methodological) 57, 289–300.

Benton, R., Vannice, K.S., Gomez-Diaz, C., and Vosshall, L.B. (2009). Variant ionotropic glutamate receptors as chemosensory receptors in Drosophila. Cell 136, 149–162.

Boucher, P., Gotthardt, M., Li, W.-P., Anderson, R.G.W., and Herz, J. (2003). LRP: role in vascular wall integrity and protection from atherosclerosis. Science 300, 329–332.

Brankatschk, M., Dunst, S., Nemetschke, L., and Eaton, S. (2014). Delivery of circulating lipoproteins to specific neurons in the Drosophila brain regulates systemic insulin signaling. Elife 3.

Carlyle, B.C., Kitchen, R.R., Kanyo, J.E., Voss, E.Z., Pletikos, M., Sousa, A.M.M., Lam, T.T., Gerstein, M.B., Sestan, N., and Nairn, A.C. (2017). A multiregional proteomic survey of the postnatal human brain. Nat. Neurosci. 20, 1787–1795.

Chen, W., and Hing, H. (2008). The L1-CAM, Neuroglian, functions in glial cells for Drosophila antennal lobe development. Dev. Neurobiol. 68, 1029–1045.

Chotard, C., Leung, W., and Salecker, I. (2005). glial cells missing and gcm2 cell autonomously regulate both glial and neuronal development in the visual system of Drosophila. Neuron 48, 237–251.

Christopoulos, A. (2002). Allosteric binding sites on cell-surface receptors: novel targets for drug discovery. Nat. Rev. Drug Discov. 1, 198–210.

Cordwell, S.J., and Thingholm, T.E. (2010). Technologies for plasma membrane proteomics. Proteomics 10, 611–627.

Couto, A., Alenius, M., and Dickson, B.J. (2005). Molecular, anatomical, and functional organization of the Drosophila olfactory system. Curr. Biol. 15, 1535–1547.

de Wit, J., Hong, W., Luo, L., and Ghosh, A. (2011). Role of leucine-rich repeat proteins in the development and function of neural circuits. Annu. Rev. Cell Dev. Biol. 27, 697–729.

Dietzl, G., Chen, D., Schnorrer, F., Su, K.-C., Barinova, Y., Fellner, M., Gasser, B., Kinsey, K., Oppel, S., Scheiblauer, S., et al. (2007). A genome-wide transgenic RNAi library for conditional gene inactivation in Drosophila. Nature 448, 151–156.

Dobin, A., Davis, C.A., Schlesinger, F., Drenkow, J., Zaleski, C., Jha, S., Batut, P., Chaisson, M., and Gingeras, T.R. (2013). STAR: ultrafast universal RNA-seq aligner. Bioinformatics 29, 15–21.

Duffy, J.B., Harrison, D.A., and Perrimon, N. (1998). Identifying loci required for follicular patterning using directed mosaics. Development 125, 2263–2271.

Fecher, C., Trovò, L., Müller, S.A., Snaidero, N., Wettmarshausen, J., Heink, S., Ortiz, O., Wagner, I., Kühn, R., Hartmann, J., et al. (2019). Cell-type-specific profiling of brain mitochondria reveals functional and molecular diversity. Nat. Neurosci. 22, 1731–1742.

Fishilevich, E., and Vosshall, L.B. (2005). Genetic and functional subdivision of the Drosophila antennal lobe. Curr. Biol. 15, 1548–1553.

Gao, Q., Yuan, B., and Chess, A. (2000). Convergent projections of Drosophila olfactory neurons to specific glomeruli in the antennal lobe. Nat. Neurosci. 3, 780–785.

Golic, K.G., and Lindquist, S. (1989). The FLP recombinase of yeast catalyzes site-specific recombination in the Drosophila genome. Cell 59, 499–509.

Gratz, S.J., Cummings, A.M., Nguyen, J.N., Hamm, D.C., Donohue, L.K., Harrison, M.M., Wildonger, J., and O’Connor-Giles, K.M. (2013). Genome engineering of Drosophila with the CRISPR RNA-guided Cas9 nuclease. Genetics 194, 1029–1035.

Gratz, S.J., Ukken, F.P., Rubinstein, C.D., Thiede, G., Donohue, L.K., Cummings, A.M., and O’Connor-Giles, K.M. (2014). Highly specific and efficient CRISPR/Cas9-catalyzed homology-directed repair in Drosophila. Genetics 196, 961–971.

Guajardo, R., Luginbuhl, D.J., Han, S., Luo, L., and Li, J. (2019). Functional divergence of Plexin B structural motifs in distinct steps of Drosophila olfactory circuit assembly. Elife 8.

Han, C., Jan, L.Y., and Jan, Y.-N. (2011). Enhancer-driven membrane markers for analysis of nonautonomous mechanisms reveal neuron-glia interactions in Drosophila. Proc. Natl. Acad. Sci. USA 108, 9673–9678.

Han, S., Li, J., and Ting, A.Y. (2018). Proximity labeling: spatially resolved proteomic mapping for neurobiology. Curr. Opin. Neurobiol. 50, 17–23.

Herz, J., and Strickland, D.K. (2001). LRP: a multifunctional scavenger and signaling receptor. J. Clin. Invest. 108, 779–784.

Hong, W., and Luo, L. (2014). Genetic control of wiring specificity in the fly olfactory system. Genetics 196, 17–29.

Hong, W., Zhu, H., Potter, C.J., Barsh, G., Kurusu, M., Zinn, K., and Luo, L. (2009). Leucine-rich repeat transmembrane proteins instruct discrete dendrite targeting in an olfactory map. Nat. Neurosci. 12, 1542–1550.

Hong, W., Mosca, T.J., and Luo, L. (2012). Teneurins instruct synaptic partner matching in an olfactory map. Nature 484, 201–207.

Hosp, F., and Mann, M. (2017). A primer on concepts and applications of proteomics in neuroscience. Neuron 96, 558–571.

Hummel, T., and Zipursky, S.L. (2004). Afferent induction of olfactory glomeruli requires N-cadherin. Neuron 42, 77–88.

Hummel, T., Vasconcelos, M.L., Clemens, J.C., Fishilevich, Y., Vosshall, L.B., and Zipursky, S.L. (2003). Axonal targeting of olfactory receptor neurons in Drosophila is controlled by Dscam. Neuron 37, 221–231.

Hung, V., Zou, P., Rhee, H.-W., Udeshi, N.D., Cracan, V., Svinkina, T., Carr, S.A., Mootha, V.K., and Ting, A.Y. (2014). Proteomic mapping of the human mitochondrial intermembrane space in live cells via ratiometric APEX tagging. Mol. Cell 55, 332–341.

Ito, K., Suzuki, K., Estes, P., Ramaswami, M., Yamamoto, D., and Strausfeld, N.J. (1998). The organization of extrinsic neurons and their implications in the functional roles of the mushroom bodies in Drosophila melanogaster Meigen. Learn. Mem. 5, 52–77.

Jan, Y.-N., and Jan, L.Y. (2010). Branching out: mechanisms of dendritic arborization. Nat. Rev. Neurosci. 11, 316–328.

Jefferis, G.S., Marin, E.C., Stocker, R.F., and Luo, L. (2001). Target neuron prespecification in the olfactory map of Drosophila. Nature 414, 204–208.

Jefferis, G.S.X.E., Vyas, R.M., Berdnik, D., Ramaekers, A., Stocker, R.F., Tanaka, N.K., Ito, K., and Luo, L. (2004). Developmental origin of wiring specificity in the olfactory system of Drosophila. Development 131, 117–130.

Jenett, A., Rubin, G.M., Ngo, T.-T.B., Shepherd, D., Murphy, C., Dionne, H., Pfeiffer, B.D., Cavallaro, A., Hall, D., Jeter, J., et al. (2012). A GAL4-driver line resource for Drosophila neurobiology. Cell Rep. 2, 991–1001.

Jhaveri, D., Saharan, S., Sen, A., and Rodrigues, V. (2004). Positioning sensory terminals in the olfactory lobe of Drosophila by Robo signaling. Development 131, 1903–1912.

Joo, W.J., Sweeney, L.B., Liang, L., and Luo, L. (2013). Linking cell fate, trajectory choice, and target selection: genetic analysis of Sema-2b in olfactory axon targeting. Neuron 78, 673–686.

Kanekiyo, T., Cirrito, J.R., Liu, C.-C., Shinohara, M., Li, J., Schuler, D.R., Shinohara, M., Holtzman, D.M., and Bu, G. (2013). Neuronal clearance of amyloid-β by endocytic receptor LRP1. J. Neurosci. 33, 19276–19283.

Kang, D.E., Pietrzik, C.U., Baum, L., Chevallier, N., Merriam, D.E., Kounnas, M.Z., Wagner, S.L., Troncoso, J.C., Kawas, C.H., Katzman, R., et al. (2000). Modulation of amyloid beta-protein clearance and Alzheimer’s disease susceptibility by the LDL receptor-related protein pathway. J. Clin. Invest. 106, 1159–1166.

Kaur, R., Surala, M., Hoger, S., Grössmann, N., Grimm, A., Timaeus, L., Kallina, W., and Hummel, T. (2019). Pioneer interneurons instruct bilaterality in the Drosophila olfactory sensory map. Sci. Adv. 5, eaaw5537.

Khaliullina, H., Panáková, D., Eugster, C., Riedel, F., Carvalho, M., and Eaton, S. (2009). Patched regulates Smoothened trafficking using lipoprotein-derived lipids. Development 136, 4111–4121.

Kolodkin, A.L., and Tessier-Lavigne, M. (2011). Mechanisms and molecules of neuronal wiring: a primer. Cold Spring Harb. Perspect. Biol. 3.

Komiyama, T., Sweeney, L.B., Schuldiner, O., Garcia, K.C., and Luo, L. (2007). Graded expression of semaphorin-1a cell-autonomously directs dendritic targeting of olfactory projection neurons. Cell 128, 399–410.

Landowski, L.M., Pavez, M., Brown, L.S., Gasperini, R., Taylor, B.V., West, A.K., and Foa, L. (2016). Low-density Lipoprotein Receptor-related Proteins in a Novel Mechanism of Axon Guidance and Peripheral Nerve Regeneration. J. Biol. Chem. 291, 1092–1102.

Larsen, C.W., Hirst, E., Alexandre, C., and Vincent, J.-P. (2003). Segment boundary formation in Drosophila embryos. Development 130, 5625–5635.

Lee, T., and Luo, L. (1999). Mosaic analysis with a repressible cell marker for studies of gene function in neuronal morphogenesis. Neuron 22, 451–461.

Li, H., Horns, F., Wu, B., Xie, Q., Li, J., Li, T., Luginbuhl, D.J., Quake, S.R., and Luo, L. (2017). Classifying Drosophila Olfactory Projection Neuron Subtypes by Single-Cell RNA Sequencing. Cell 171, 1206–1220.e22.

Li, H., Li, T., Horns, F., Li, J., Xie, Q., Xu, C., Wu, B., Kebschull, J.M., Vacek, D., Xie, A., et al. (2019). Coordinating receptor expression and wiring specificity in olfactory receptor neurons. BioRxiv. https://doi.org/10.1101/594895

Li, J., Guajardo, R., Xu, C., Wu, B., Li, H., Li, T., Luginbuhl, D.J., Xie, X., and Luo, L. (2018). Stepwise wiring of the Drosophila olfactory map requires specific Plexin B levels. Elife 7:e39088.

Lin, D.M., and Goodman, C.S. (1994). Ectopic and increased expression of Fasciclin II alters motoneuron growth cone guidance. Neuron 13, 507–523.

Liu, C.-C., Hu, J., Zhao, N., Wang, J., Wang, N., Cirrito, J.R., Kanekiyo, T., Holtzman, D.M., and Bu, G. (2017). Astrocytic LRP1 mediates brain aβ clearance and impacts amyloid deposition. J. Neurosci. 37, 4023–4031.

Liu, H.-H., McClatchy, D.B., Schiapparelli, L., Shen, W., Yates, J.R., and Cline, H.T. (2018). Role of the visual experience-dependent nascent proteome in neuronal plasticity. Elife 7.

Loh, K.H., Stawski, P.S., Draycott, A.S., Udeshi, N.D., Lehrman, E.K., Wilton, D.K., Svinkina, T., Deerinck, T.J., Ellisman, M.H., Stevens, B., et al. (2016). Proteomic analysis of unbounded cellular compartments: synaptic clefts. Cell 166, 1295–1307.e21.

Malenka, R.C., and Bear, M.F. (2004). LTP and LTD: an embarrassment of riches. Neuron 44, 5–21.

May, P., Rohlmann, A., Bock, H.H., Zurhove, K., Marth, J.D., Schomburg, E.D., Noebels, J.L., Beffert, U., Sweatt, J.D., Weeber, E.J., et al. (2004). Neuronal LRP1 functionally associates with postsynaptic proteins and is required for normal motor function in mice. Mol. Cell. Biol. 24, 8872–8883.

Mosca, T.J., and Luo, L. (2014). Synaptic organization of the Drosophila antennal lobe and its regulation by the Teneurins. Elife 3, e03726.

Natividad, L.A., Buczynski, M.W., McClatchy, D.B., and Yates, J.R. (2018). From synapse to function: A perspective on the role of neuroproteomics in elucidating mechanisms of drug addiction. Proteomes 6.

Ni, J.-Q., Zhou, R., Czech, B., Liu, L.-P., Holderbaum, L., Yang-Zhou, D., Shim, H.-S., Tao, R., Handler, D., Karpowicz, P., et al. (2011). A genome-scale shRNA resource for transgenic RNAi in Drosophila. Nat. Methods 8, 405–407.

Nunomura, K., Nagano, K., Itagaki, C., Taoka, M., Okamura, N., Yamauchi, Y., Sugano, S., Takahashi, N., Izumi, T., and Isobe, T. (2005). Cell surface labeling and mass spectrometry reveal diversity of cell surface markers and signaling molecules expressed in undifferentiated mouse embryonic stem cells. Mol. Cell Proteomics 4, 1968–1976.

Perkins, L.A., Holderbaum, L., Tao, R., Hu, Y., Sopko, R., McCall, K., Yang-Zhou, D., Flockhart, I., Binari, R., Shim, H.-S., et al. (2015). The transgenic rnai project at harvard medical school: resources and validation. Genetics 201, 843–852.

Picelli, S., Faridani, O.R., Björklund, A.K., Winberg, G., Sagasser, S., and Sandberg, R. (2014). Full-length RNA-seq from single cells using Smart-seq2. Nat. Protoc. 9, 171–181.

Port, F., Chen, H.-M., Lee, T., and Bullock, S.L. (2014). Optimized CRISPR/Cas tools for efficient germline and somatic genome engineering in Drosophila. Proc. Natl. Acad. Sci. USA 111, E2967–76.

Ritchie, M.E., Phipson, B., Wu, D., Hu, Y., Law, C.W., Shi, W., and Smyth, G.K. (2015). limma powers differential expression analyses for RNA-sequencing and microarray studies. Nucleic Acids Res. 43, e47.

Rohlmann, A., Gotthardt, M., Hammer, R.E., and Herz, J. (1998). Inducible inactivation of hepatic LRP gene by cre-mediated recombination confirms role of LRP in clearance of chylomicron remnants. J. Clin. Invest. 101, 689–695.

Shen, H.-C., Chu, S.-Y., Hsu, T.-C., Wang, C.-H., Lin, I.-Y., and Yu, H.-H. (2017). Semaphorin-1a prevents Drosophila olfactory projection neuron dendrites from mis-targeting into select antennal lobe regions. PLoS Genet. 13, e1006751.

Silbering, A.F., Rytz, R., Grosjean, Y., Abuin, L., Ramdya, P., Jefferis, G.S.X.E., and Benton, R. (2011). Complementary function and integrated wiring of the evolutionarily distinct Drosophila olfactory subsystems. J. Neurosci. 31, 13357–13375.

Singh, A.P., VijayRaghavan, K., and Rodrigues, V. (2010). Dendritic refinement of an identified neuron in the Drosophila CNS is regulated by neuronal activity and Wnt signaling. Development 137, 1351–1360.

Sperry, R.W. (1963). CHEMOAFFINITY IN THE ORDERLY GROWTH OF NERVE FIBER PATTERNS AND CONNECTIONS. Proc. Natl. Acad. Sci. USA 50, 703–710.

Sweeney, L.B., Couto, A., Chou, Y.-H., Berdnik, D., Dickson, B.J., Luo, L., and Komiyama, T. (2007). Temporal target restriction of olfactory receptor neurons by Semaphorin-1a/PlexinA-mediated axon-axon interactions. Neuron 53, 185–200.

Sweeney, L.B., Chou, Y.-H., Wu, Z., Joo, W., Komiyama, T., Potter, C.J., Kolodkin, A.L., Garcia, K.C., and Luo, L. (2011). Secreted semaphorins from degenerating larval ORN axons direct adult projection neuron dendrite targeting. Neuron 72, 734–747.

Tachibana, M., Holm, M.-L., Liu, C.-C., Shinohara, M., Aikawa, T., Oue, H., Yamazaki, Y., Martens, Y.A., Murray, M.E., Sullivan, P.M., et al. (2019). *APOE4*-mediated amyloid-β pathology depends on its neuronal receptor LRP1. J. Clin. Invest.

Tan, D.J.L., Dvinge, H., Christoforou, A., Bertone, P., Martinez Arias, A., and Lilley, K.S. (2009). Mapping organelle proteins and protein complexes in Drosophila melanogaster. J. Proteome Res. 8, 2667–2678.

Thompson, A., Schäfer, J., Kuhn, K., Kienle, S., Schwarz, J., Schmidt, G., Neumann, T., Johnstone, R., Mohammed, A.K.A., and Hamon, C. (2003). Tandem mass tags: a novel quantification strategy for comparative analysis of complex protein mixtures by MS/MS. Anal. Chem. 75, 1895–1904.

Tirian, L., and Dickson, B. (2017). The VT GAL4, LexA, and split-GAL4 driver line collections for targeted expression in the Drosophila nervous system. BioRxiv.

Vosshall, L.B., and Stocker, R.F. (2007). Molecular architecture of smell and taste in Drosophila. Annu. Rev. Neurosci. 30, 505–533.

Vosshall, L.B., Wong, A.M., and Axel, R. (2000). An olfactory sensory map in the fly brain. Cell 102, 147–159.

Ward, A., Hong, W., Favaloro, V., and Luo, L. (2015). Toll receptors instruct axon and dendrite targeting and participate in synaptic partner matching in a Drosophila olfactory circuit. Neuron 85, 1013–1028.

Watts, R.J., Schuldiner, O., Perrino, J., Larsen, C., and Luo, L. (2004). Glia engulf degenerating axons during developmental axon pruning. Curr. Biol. 14, 678–684.

Wilson, R.I. (2013). Early olfactory processing in Drosophila: mechanisms and principles. Annu. Rev. Neurosci. 36, 217–241.

Wollscheid, B., Bausch-Fluck, D., Henderson, C., O’Brien, R., Bibel, M., Schiess, R., Aebersold, R., and Watts, J.D. (2009). Mass-spectrometric identification and relative quantification of N-linked cell surface glycoproteins. Nat. Biotechnol. 27, 378–386.

Wu, J.S., and Luo, L. (2006a). A protocol for mosaic analysis with a repressible cell marker (MARCM) in Drosophila. Nat. Protoc. 1, 2583–2589.

Wu, J.S., and Luo, L. (2006b). A protocol for dissecting Drosophila melanogaster brains for live imaging or immunostaining. Nat. Protoc. 1, 2110–2115.

Xie, Q., Wu, B., Li, J., Xu, C., Li, H., Luginbuhl, D.J., Wang, X., Ward, A., and Luo, L. (2019). Transsynaptic Fish-lips signaling prevents misconnections between nonsynaptic partner olfactory neurons. Proc. Natl. Acad. Sci. USA 116.32 (2019): 16068–16073.

Xu, T., and Rubin, G.M. (1993). Analysis of genetic mosaics in developing and adult Drosophila tissues. Development 117, 1223–1237.

Yu, H.-H., Kao, C.-F., He, Y., Ding, P., Kao, J.-C., and Lee, T. (2010). A complete developmental sequence of a Drosophila neuronal lineage as revealed by twin-spot MARCM. PLoS Biol. 8.

Zhang, W., Zhou, G., Zhao, Y., White, M.A., and Zhao, Y. (2003). Affinity enrichment of plasma membrane for proteomics analysis. Electrophoresis 24, 2855–2863.

Zhu, H., and Luo, L. (2004). Diverse functions of N-cadherin in dendritic and axonal terminal arborization of olfactory projection neurons. Neuron 42, 63–75.

Zhu, H., Hummel, T., Clemens, J.C., Berdnik, D., Zipursky, S.L., and Luo, L. (2006). Dendritic patterning by Dscam and synaptic partner matching in the Drosophila antennal lobe. Nat. Neurosci. 9, 349–355.

Zipursky, S.L., and Sanes, J.R. (2010). Chemoaffinity revisited: dscams, protocadherins, and neural circuit assembly. Cell 143, 343–353.

